# Competitive elimination of ZO-1/ZO-2-deficient cells regulates epithelial barrier homeostasis

**DOI:** 10.1101/2025.01.10.632304

**Authors:** Tetsuhisa Otani, Thanh Phuong Nguyen, Noriyuki Kinoshita, Toshihiko Fujimori, Mikio Furuse

## Abstract

Epithelia cover the body and form a barrier to segregate the internal body from the external environment. Epithelial tissues contact the external environment and are exposed to various stresses that are potentially deleterious for the epithelial barrier^1^. However, how focal epithelial barrier defects are detected and repaired to maintain epithelial barrier homeostasis remains poorly understood. We co-cultured *ZO-1*/*ZO-2* double-knockout (DKO) MDCK II cells, which lack tight junctions, with wild-type MDCK II cells, to understand how epithelial cells respond to focal epithelial barrier defects. When co-cultured with wild-type cells, *ZO-1*/*ZO-2* DKO cells were selectively eliminated by induction of apoptosis. The elimination depended on a purse-string-like contraction of supracellular actomyosin cables formed at the clone boundary, regulated by ROCK. Furthermore, Hippo signaling and adherens junctions in the surrounding wild-type cells were required to eliminate *ZO-1*/*ZO-2* DKO cells. These results demonstrate that selective elimination of ZO-1/ZO-2-deficient cells by cell competition regulates epithelial barrier homeostasis.

**In brief:** Otani et al. reveal that epithelial barrier homeostasis is regulated by cell competition-mediated elimination of ZO-1/ZO-2-deficient cells. Supracellular actomyosin cables form at the clone boundary and constrict in a purse-string-like manner to eliminate the ‘loser’ cells. Mechanosensing in the winning cells is required to eliminate the ‘loser’ cells.

**Highlights:**

- ZO-1/ZO-2-deficient cells are eliminated by cell competition
- Actomyosin cables form in the winning cells at the clone boundary
- A purse-string-like contraction of actomyosin cables compresses the losing cells
- Mechanosensing in the winning cells is important to eliminate the ‘loser’ cells

## Introduction

Epithelial cells form a sheet surrounding the body as a barrier to protect the internal body from the external environment. To form an epithelial barrier, it is essential to seal the intercellular space to restrict the paracellular diffusion of solutes. Epithelial cells form intercellular junctions to adhere together, and tight junctions (TJs), adherens junctions (AJs), and desmosomes are collectively termed the apical junctional complex^2^. TJs are located at the most apical region of the junctional complex, and are responsible for occluding the intercellular space to form the epithelial barrier, while AJs interact with the circumferential actomyosin ring and play important roles in intercellular force transmission^3,4^. Integral membrane proteins, including claudins, occludin, and immunoglobulin superfamily proteins, localize to TJs and interact with zonula occludens (ZO) family scaffolding proteins through their cytoplasmic region^4,5^. ZO-1 and ZO-2 are essential for TJ assembly, and cells that lack ZO-1 and ZO-2 exhibit pleiotropic phenotypes, including loss of TJ structure, disruption of the epithelial barrier, epithelial polarity defects, and actomyosin and AJ disorganization^6–9^. AJs contain the membrane proteins cadherins and nectins, which interact with the actin cytoskeleton via the linker proteins β-/α-catenin or afadin^10^. α-catenin can undergo a tension-dependent conformation change and act as a mechanosensor^11^.

Epithelia are located at the surface of the body and directly face the external environment. They are exposed to various stresses, including infection and mechanical, chemical, and thermal stress, which challenge TJ function^1^. For example, epithelial barrier function and TJ organization are compromised by infection and other stress^12^. Such epithelial barrier defects challenge the function of the body and are associated with various diseases^5^. It is thus reasonable to assume that mechanisms exist to prevent, detect, and repair epithelial barrier defects. However, it is poorly understood how homeostasis of the epithelial barrier is maintained.

Previous studies have revealed several mechanisms involved in epithelial barrier homeostasis. For example, TJs are required for epithelial junctions to resist mechanical forces and prevent their mechanical rupture^13^, and *de novo* TJ assembly accompanies cytokinesis or cell extrusion to maintain the epithelial barrier function during epithelial cell turnover^14–18^. Micrometer-scale TJ disruptions are repaired by Rho-dependent actin assembly and myosin contraction or by the release of reservoir claudin molecules^19–21^, and macroscopic TJ dysfunctions are repaired by TJ reassembly induced by junction-inducing peptides during inflammation^22^. However, how epithelial cells cope with mesoscopic focal epithelial barrier defects has not been clarified.

Cell competition is a mechanism that eliminates unfit cells from tissues^23,24^. Cell competition was originally characterized in *Drosophila*^25^, and regulates tissue homeostasis by eliminating oncogenic cells, developmentally delayed or misspecified cells, and unfit stem cells^26–34^. However, what triggers such cell competition, how the losing cells are eliminated, and the physiological roles of cell competition remain poorly characterized.

Here, we demonstrate that ZO-1/ZO-2-deficient cells are eliminated from epithelia by cell competition. Moreover, the interfacial tension generated by supracellular actomyosin cables formed at the boundary between the winning and losing cells is crucial to eliminate ZO-1/ZO-2-deficient cells. These findings suggest that cell competition regulates epithelial barrier homeostasis.

## Results

### *ZO-1/ZO-2* double-knockout cells are eliminated by cell competition

To understand how epithelia respond to focal epithelial barrier defects, we established a co-culture system in which we cultured parental MDCK II cells together at a 1:1 ratio with *ZO-1/ZO-2* double-knockout (DKO) cells that lack TJs (Fig. 1A), and examined the epithelial barrier function by transepithelial electric resistance (TER) measurements. The co-culture initially exhibited low barrier function, reflecting the presence of focal epithelial barrier defects. However, epithelial barrier function was progressively restored, and the TER values became comparable to parental MDCK II cells by 6 d (Fig. 1B). Staining of the co-culture with ZO-1 revealed that ZO-1-positive parental cells and ZO-1-negative DKO cells initially mixed and formed a continuous monolayer (Fig. 1C). However, ZO-1-negative cells were progressively eliminated from the epithelial monolayer, and a continuous chicken-wire pattern of ZO-1 staining was restored by 5 d (Fig. 1C). Expression of ZO-1 linked to enhanced green fluorescent protein (ZO-1-EGFP) in *ZO-1/ZO-2* DKO cells completely suppressed their elimination (Fig. 1D), and *ZO-1*-knockout (KO)/*ZO-2*-knockdown (KD) EpH4 cells were also eliminated upon co-culture with parental EpH4 cells (Fig. 1E). These results suggest that ZO-1/ZO-2-deficient cells are eliminated from the epithelial monolayer upon co-culture with wild-type (WT) cells.

**Figure 1.**
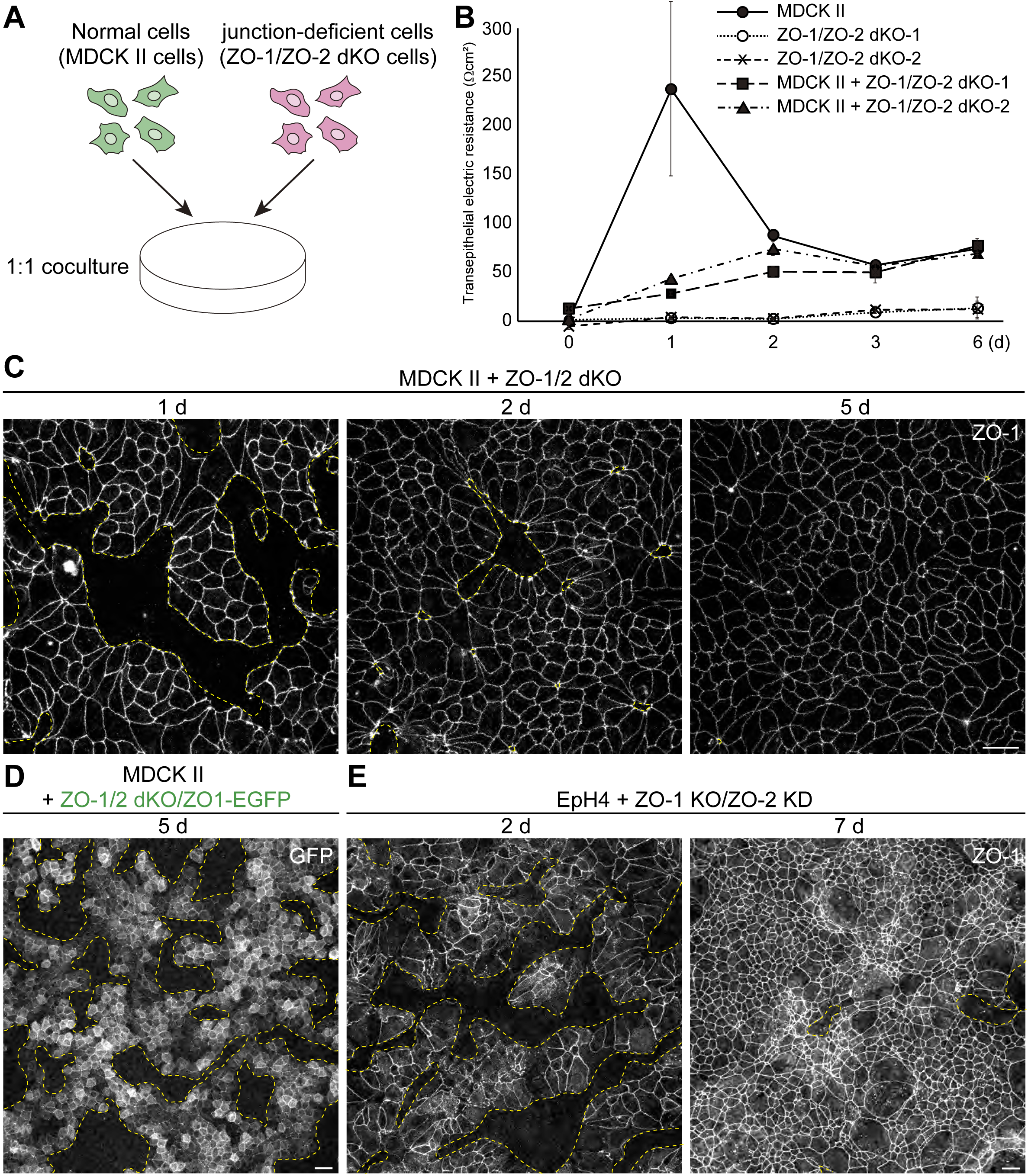
*ZO-1*/*ZO-2* DKO cells are eliminated by cell competition. (A) Schematic illustration of the experimental design. Normal MDCK II cells and *ZO-1*/*ZO-2* DKO cells were co-cultured at a 1:1 ratio. (B) Temporal changes of TER values in the MDCK II or *ZO-1*/*ZO-2* DKO solo cultures versus MDCK II and *ZO-1*/*ZO-2* DKO co-culture. The data represent the TER values and are presented as mean ± SD (*n* = 3 each). The TER values of the co-culture became comparable to the WT MDCK II cells by 3 d after co-culture. (C) ZO-1 staining of the co-culture between WT MDCK II cells and *ZO-1*/*ZO-2* DKO cells. ZO-1-negative *ZO-1*/*ZO-2* DKO cells were progressively eliminated. Clone boundaries are indicated by yellow dotted lines throughout the manuscript. (D) Elimination of *ZO-1*/*ZO-2* DKO cells was suppressed by re-expression of ZO-1-EGFP. (E) Elimination of *ZO-1* KO/*ZO-2* KD EpH4 cells upon co-culture with normal EpH4 cells. Scale bars: 20 μm.

### *ZO-1*/*ZO-2* DKO cells are eliminated by induction of apoptosis

To clarify how *ZO-1*/*ZO-2* DKO cells are eliminated, we expressed EGFP in *ZO-1*/*ZO-2* DKO cells and co-cultured them with parental MDCK II cells. Time-lapse imaging revealed that the *ZO-1*/*ZO-2* DKO cells progressively underwent cell death when co-cultured with parental MDCK II cells (Fig. 2A, B, Video S1), resulting in a reduction of the area occupied by *ZO-1*/*ZO-2* DKO cells (Fig. 2B). *ZO-1*/*ZO-2* DKO cells fragmented upon cell death, suggesting that they underwent apoptosis. Cleaved caspase-3 antibody staining or caspase-3/7 *in situ* zymography revealed that caspase is activated in *ZO-1*/*ZO-2* DKO cells upon co-culture with parental MDCK II cells (Fig. 2C).

**Figure 2.**
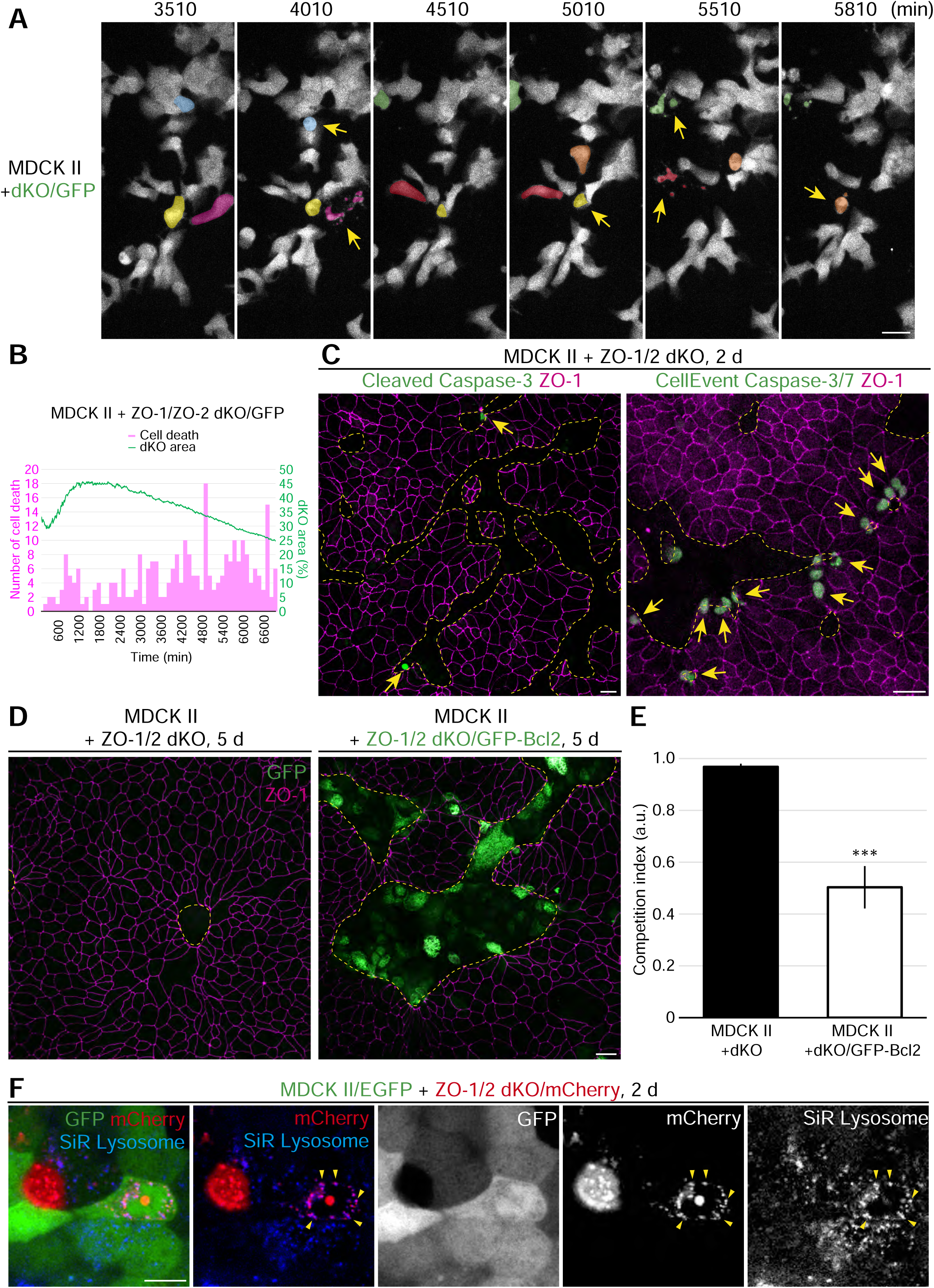
***ZO-1*/*ZO-2* DKO cells undergo apoptosis when surrounded by WT cells.** (A) Snapshots from time-lapse imaging of MDCK II cells co-cultured with *ZO-1*/*ZO-2* DKO cells expressing EGFP. EGFP-positive *ZO-1*/*ZO-2* DKO cells progressively underwent cell death (yellow arrow). Dying cells are pseudo-colored. (B) Quantification of the numbers of cell deaths (magenta) and the area occupied by *ZO-1*/*ZO-2* DKO cells (green). *ZO-1*/*ZO-2* DKO cells were progressively eliminated. (C) Caspase activation in *ZO-1*/*ZO-2* DKO cells upon co-culture with MDCK II cells. Caspase-3 activity (yellow arrow) was visualized by anti-cleaved caspase-3 antibody staining (left, green) or caspase-3/7 *in situ* zymography by CellEvent Caspase-3/7 (right, green), and co-stained with anti-ZO-1 antibodies (magenta). Note that some cells are apically extruded. (D) Overexpression of an apoptosis inhibitor, GFP-BCL2 (green), in *ZO-1*/*ZO-2* DKO cells suppressed their elimination. *ZO-1*/*ZO-2* DKO cells expressing GFP-BCL2 were co-cultured with MDCK II cells and co-stained with anti-GFP (green) and anti-ZO-1 antibodies (magenta). (E) The efficiency of the elimination of *ZO-1*/*ZO-2* DKO cells was quantified by measuring the competition index, which is given by (A^MDCK^ ^II^ – A^KO^) / (A^MDCK^ ^II^ + A^KO^), where A^MDCK^ ^II^ is the area occupied by surrounding WT cells, and A^KO^ is the area occupied by *ZO-1*/*ZO-2* DKO cells. Overexpression of GFP-BCL2 significantly suppressed the elimination of *ZO-1*/*ZO-2* DKO cells. Data represent the mean ± SD (*n* = 5 each), ***p < 0.0005, compared by *t*-test. (F) *ZO-1*/*ZO-2* DKO cells are engulfed by surrounding MDCK II cells. MDCK II cells labeled with EGFP were co-cultured with *ZO-1*/*ZO-2* DKO cells expressing mCherry, and lysosomes were visualized by SiR-lysosome. mCherry-positive cell debris was observed within the lysosomes of EGFP-positive cells (yellow arrowheads). Scale bars: 20 μm (C, D); 10 μm (F). See also Video S1.

Moreover, overexpression of *BCL2*, an inhibitor of apoptosis^35^, suppressed the elimination of *ZO-1*/*ZO-2* DKO cells (Fig. 2D, E). When mCherry-expressing *ZO-1*/*ZO-2* DKO cells were co-cultured with EGFP-MDCK II cells, mCherry-positive cell debris was observed within EGFP-positive MDCK II cells and overlapped with lysosome markers, suggesting that some of the *ZO-1*/*ZO-2* DKO cells were engulfed by neighboring cells (Fig. 2F). Taken together, these results demonstrate that *ZO-1*/*ZO-2* DKO cells are eliminated by selective induction of apoptosis upon co-culture with WT epithelial cells.

### Actomyosin cables regulate the elimination of *ZO-1*/*ZO-2* DKO cells

To uncover the molecular mechanisms underlying the elimination of *ZO-1*/*ZO-2* DKO cells, we treated the co-culture with a chemical compound library with known targets and screened for compounds that inhibited the elimination of *ZO-1*/*ZO-2* DKO cells (Fig. 3A). Visual screening by microscopy identified several compounds that inhibited the elimination of *ZO-1*/*ZO-2* DKO cells, including inhibitors of the ROCK, mTOR, and TGFβR/ALK pathways (Fig. 3B, S1, Table S1). Reanalysis of these pathways revealed that ROCK inhibitors robustly and reproducibly inhibited the elimination of *ZO-1*/*ZO-2* DKO cells (Fig. 3C, D, S2).

**Figure 3.**
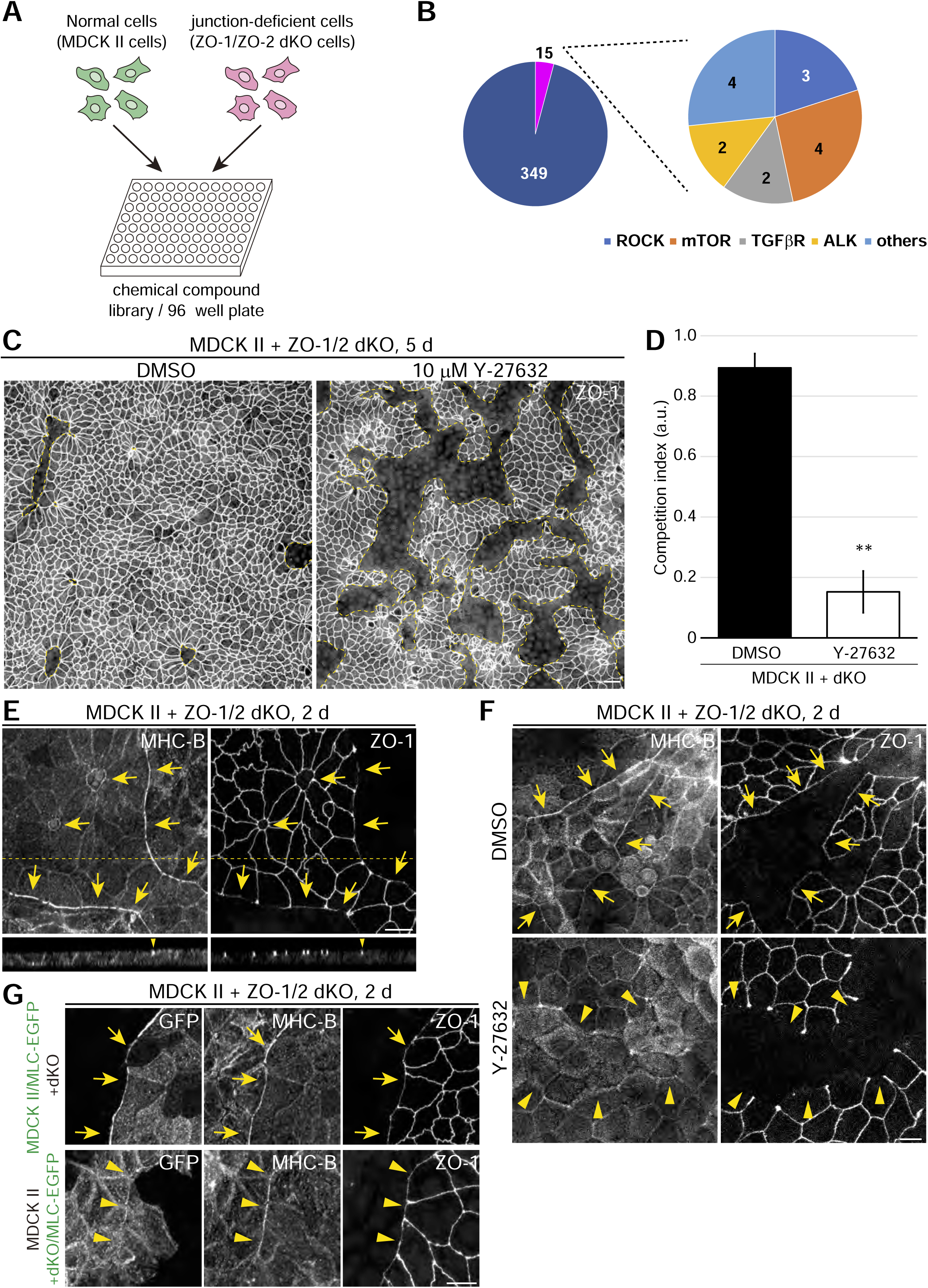
ROCK regulates the elimination of *ZO-1*/*ZO-2* DKO cells by regulating the formation of supracellular actomyosin cables at the clone boundary. (A) Schematic illustration of the chemical compound library screening. MDCK II cells and *ZO-1*/*ZO-2* DKO cells were co-cultured at a 1:1 ratio in 96-well plates containing the chemical compound library for 5 d, and subsequently stained with anti-ZO-1 antibodies. Each well was visually examined by microscopic observation. (B) Summary of the chemical compound library screening. Of the 364 compounds screened, 15 compounds qualitatively suppressed the elimination of *ZO-1*/*ZO-2* DKO cells. Several compounds targeted common pathways, including ROCK (3 compounds), mTOR (4 compounds), and TGFβR/ALK (4 compounds). (C) Effect of the ROCK inhibitor Y-27632 on the elimination of *ZO-1*/*ZO-2* DKO cells. Administration of 10 μM Y-27632 inhibited *ZO-1*/*ZO-2* DKO cell elimination. (D) Quantification of the effect of Y-27632. Data represent the mean ± SD (*n* = 3 each), **p < 0.001, compared by *t*-test. (E) Actomyosin organization was visualized by anti-Myosin IIB (MHC-B) staining. Supracellular actomyosin cables assembled at the clone boundaries between the *ZO-1*/*ZO-2* DKO cells and MDCK II cells (yellow arrows). x–z sections at the yellow dotted line show that the actomyosin cables are anchored to the apical junctions (yellow arrowhead). (F) Y-27632 treatment inhibited supracellular actomyosin cable formation at the clone boundaries. In control DMSO-treated cells, supracellular actomyosin cables were observed at the clone boundaries (yellow arrows), whereas supracellular actomyosin cable formation was suppressed in Y-27632-treated cells (yellow arrowheads). (G) The supracellular actomyosin cables formed in the surrounding WT cells. The supracellular actomyosin cables were labeled when myosin light chain-EGFP (MLC-EGFP) was expressed in MDCK II cells (yellow arrows), but not when MLC-EGFP was expressed in *ZO-1*/*ZO-2* DKO cells (yellow arrowheads). Scale bars: 20 μm (C); 10 μm (E–G). See also Figures S1–S3.

ROCK is a central regulator of actomyosin organization^36^. Myosin IIB immunostaining revealed that supracellular actomyosin cables formed at the boundary between parental MDCK II cells and *ZO-1*/*ZO-2* DKO cells upon co-culture (Fig. 3E). The actomyosin cables were formed at the AJ level, and the actomyosin cables of neighboring cells connected via the AJs (Fig. 3E). Treatment with the ROCK inhibitor Y-27632 inhibited actomyosin cable formation (Fig. 3F). To elucidate which cell population forms the supracellular actomyosin cables, we expressed MLC (myosin light chain)-EGFP or mVenus-LifeAct in MDCK II or *ZO-1*/*ZO-2* DKO cells and examined their localization. The supracellular actomyosin cables were labeled by MLC-EGFP or mVenus-LifeAct expressed in MDCK II cells, but not in *ZO-1*/*ZO-2* DKO cells (Fig. 3G, S3), suggesting that the actomyosin cables formed in MDCK II cells that neighbored *ZO-1*/*ZO-2* DKO cells. These results suggest that ROCK-dependent formation of supracellular actomyosin cables regulates the elimination of *ZO-1*/*ZO-2* DKO cells.

### *ZO-1*/*ZO-2* DKO cells are eliminated by purse-string-like contractions of the actomyosin cable

To elucidate how the supracellular actomyosin cables regulate the elimination of *ZO-1*/*ZO-2* DKO cells, we performed time-lapse imaging of MDCK II cells expressing MLC-EGFP co-cultured with *ZO-1*/*ZO-2* DKO cells. Intriguingly, when the parental MDCK II cells surrounded *ZO-1*/*ZO-2* DKO cells, the supracellular actomyosin cables contracted in a purse-string-like manner and eliminated the *ZO-1*/*ZO-2* DKO cells (Fig. 4A, Video S2). This raises the possibility that the *ZO-1*/*ZO-2* DKO cells are eliminated by mechanical cell competition driven by the purse-string-like contraction of supracellular actomyosin cables surrounding the losing cells.

**Figure 4.**
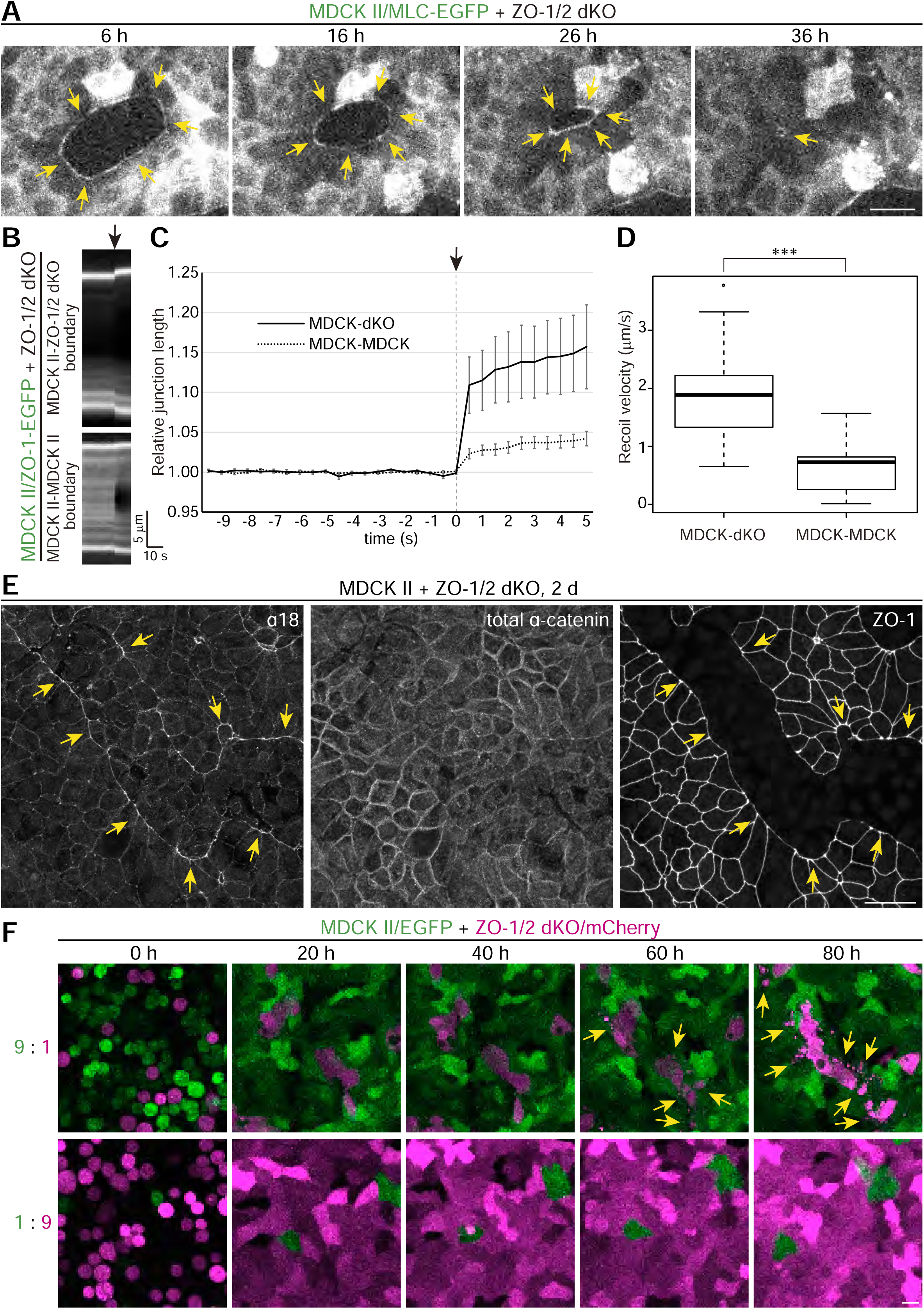
Purse-string-like contraction of the actomyosin cables eliminates *ZO-1*/*ZO-2* DKO cells. (A) Snapshots from time-lapse imaging of MDCK II cells expressing MLC-EGFP co-cultured with *ZO-1*/*ZO-2* DKO cells. Supracellular actomyosin cables (yellow arrows) form and surround the *ZO-1*/*ZO-2* DKO cells and undergo a purse-string-like contraction. (B) Representative kymographs of laser ablation experiments of MDCK II cells expressing ZO-1-EGFP co-cultured with *ZO-1*/*ZO-2* DKO cells. Cell junctions between MDCK II cells and junctions between MDCK II and *ZO-1*/*ZO-2* DKO cells were ablated. (C) Time course of cell junction recoiling on laser ablation. Black arrows in B and C indicate the time point when laser ablation was performed. Data are presented as mean ± SD (*n* = 17 for MDCK–DKO boundaries, *n* = 15 for MDCK– MDCK boundaries). (D) Initial recoil velocity of the cell junctions upon laser ablation. The junctions between MDCK II and *ZO-1*/*ZO-2* DKO cells showed higher recoil velocities, demonstrating that the clone boundaries are under higher tension. ***p < 0.00005, compared by *t*-test. (E) Immunostaining by α18 antibodies, which recognize a tension-sensitive epitope of α-catenin, together with total α-catenin and ZO-1 staining. The α18 epitope was enriched at the clonal boundaries (yellow arrows), while the total α-catenin was evenly distributed along cell–cell junctions, suggesting that the clonal boundaries are under higher tension. (F) Snapshots from time-lapse imaging of MDCK II cells expressing EGFP co-cultured with *ZO-1*/*ZO-2* DKO cells expressing mCherry. *ZO-1*/*ZO-2* DKO cells were progressively eliminated when surrounded by MDCK II cells (yellow arrows, 9:1 co-culture), while elimination did not proceed when *ZO-1*/*ZO-2* DKO cells surrounded MDCK II cells (1:9 co-culture). Scale bars: 20 μm. See also Figure S4 and Videos S2–S4.

To test whether junctional tension is increased at the clonal boundaries, we performed laser ablation of the cell junctions in MDCK II cells expressing ZO-1-EGFP co-cultured with *ZO-1*/*ZO-2* DKO cells. Laser ablation experiments showed that the recoil velocity was significantly elevated at the clonal boundaries (0.65 ± 0.44 μm/s for MDCK II–MDCK II junctions; 1.87 ± 0.86 μm/s for MDCK II–DKO junctions) (Fig. 4B–D, Video S3). Consistently, α18 antibodies, which detect a tension-sensitive epitope of α-catenin^10^, strongly stained the clone boundary between parental MDCK II cells and *ZO-1*/*ZO-2* DKO cells, while total α-catenin was not enriched (Fig. 4E). These results indicate that the clonal boundaries are under elevated tension.

The contraction of the supracellular actomyosin cables suggests that the *ZO-1*/*ZO-2* DKO cells are compressed. Quantification of the cell morphology showed that *ZO-1*/*ZO-2* DKO cells have smaller apical areas, and that parental MDCK II cells at the clonal boundary elongate towards *ZO-1*/*ZO-2* DKO cells in a ROCK-dependent manner (Fig. S4). These results suggest that the purse-string-like contraction of the supracellular actomyosin cables compresses the *ZO-1*/*ZO-2* DKO cells and induces apoptosis.

This contraction of the supracellular actomyosin cables suggests that *ZO-1*/*ZO-2* DKO cells are eliminated by compression-induced cell death. However, an alternative possibility is that cell death is induced by a chemical signal coming from the neighboring WT cells, and the purse-string-like contraction occurs to extrude the dying cells. To distinguish these possibilities, we altered the ratio of the co-culture of MDCK II and *ZO-1*/*ZO-2* DKO cells so that the cells were co-cultured at either a 9:1 or a 1:9 ratio. If the death of *ZO-1*/*ZO-2* DKO cells is induced by a chemical signal, apoptosis would be induced regardless of the co-culture ratio. However, if mechanical compression is required to eliminate *ZO-1*/*ZO-2* DKO cells, cell death will only be induced when the *ZO-1*/*ZO-2* DKO cells are surrounded by WT cells. When the MDCK II cells and *ZO-1*/*ZO-2* DKO cells were co-cultured at a 9:1 ratio, the *ZO-1*/*ZO-2* DKO cells were surrounded by MDCK II cells and underwent apoptosis (Fig. 4F, Video S4). However, when MDCK II cells and *ZO-1*/*ZO-2* DKO cells were co-cultured at a 1:9 ratio and the *ZO-1*/*ZO-2* DKO cells were not surrounded by MDCK II cells, no cell death was observed (Fig. 4F, Video S4). These results indicate that the *ZO-1*/*ZO-2* DKO cells are eliminated by mechanical cell competition driven by the purse-string-like contraction of the supracellular actomyosin cables.

### Adherens junctions of the surrounding cells are required for the elimination of *ZO-1*/*ZO-2* DKO cells

The anchoring of the supracellular actomyosin cables to the AJ and the enrichment of the α18 epitope at the clonal boundary suggest that AJs may play important roles in the elimination of *ZO-1*/*ZO-2* DKO cells. Interestingly, afadin, an AJ-localized actin-binding protein, was strongly enriched at the clonal boundary (Fig. 5A) in a myosin II-dependent manner (Fig. 5B). To investigate whether AJs are required to eliminate *ZO-1*/*ZO-2* DKO cells, we examined the roles of E-cadherin and afadin. Knockout of E-cadherin or afadin in the surrounding WT cells inhibited the elimination of *ZO-1*/*ZO-2* DKO cells (Fig. 5C–F, Video S5). Moreover, the compression of *ZO-1*/*ZO-2* DKO cells was suppressed by knockout of E-cadherin or afadin, and the directional elongation of the neighboring WT cells was perturbed (Fig. S4). These results demonstrate that AJs are essential for the elimination of *ZO-1*/*ZO-2* DKO cells.

**Figure 5.**
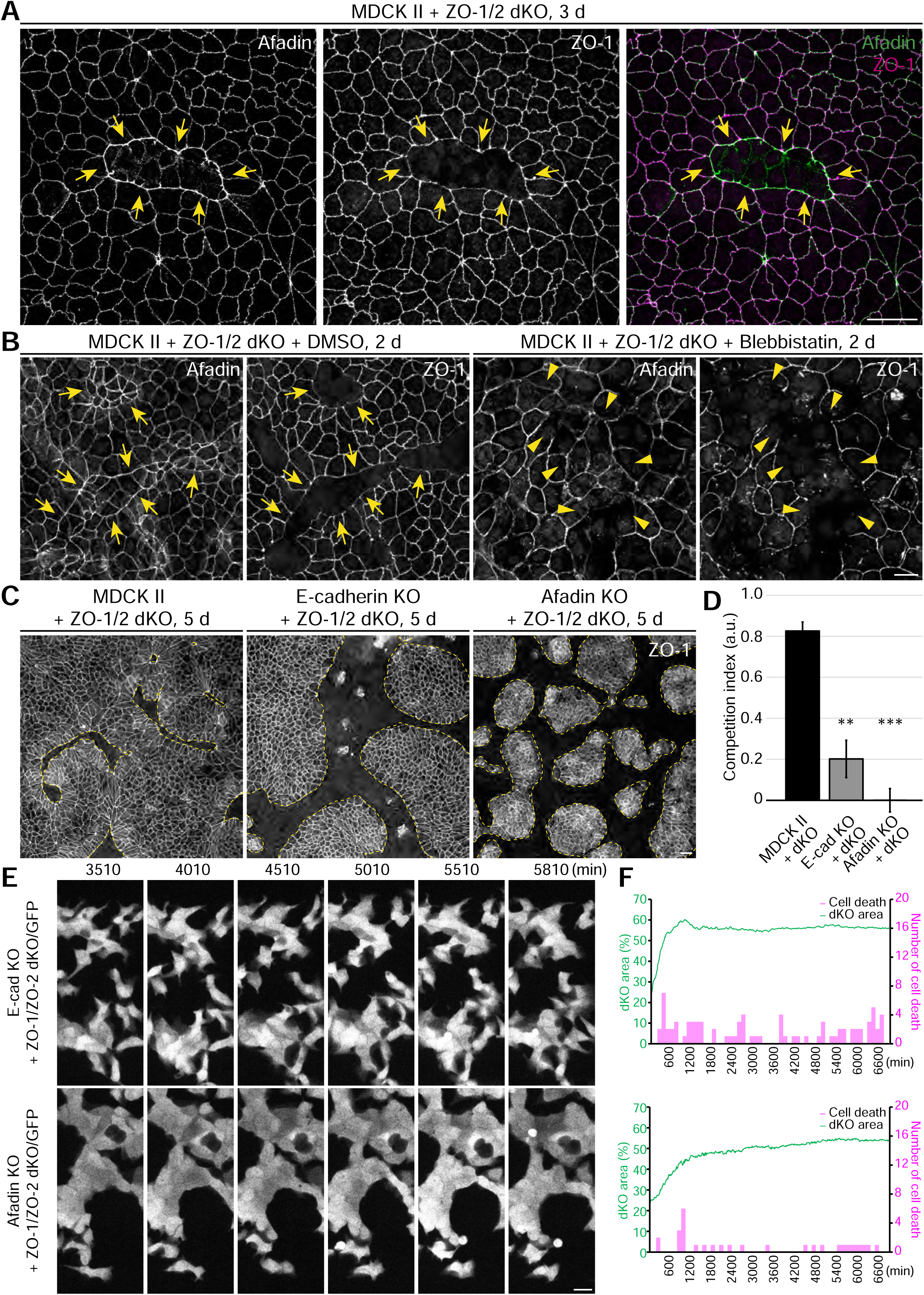
AJs in the surrounding WT cells are required to eliminate *ZO-1*/*ZO-2* DKO cells. (A) MDCK II cells and *ZO-1*/*ZO-2* DKO cells were co-cultured and stained with anti-afadin (green) and anti-ZO-1 antibodies (magenta). Afadin was enriched at the clone boundaries (yellow arrows). (B) MDCK II cells and *ZO-1*/*ZO-2* DKO cells were co-cultured in the presence of DMSO or 50 μM blebbistatin. The accumulation of afadin at the clonal boundaries (yellow arrows) was suppressed when Myosin II activity was inhibited by blebbistatin (yellow arrowheads). (C) *ZO-1*/*ZO-2* DKO cell elimination was suppressed when E-cadherin or afadin was knocked out in the surrounding MDCK II cells. (D) Quantification of the co-culture of MDCK II, *E-cadherin* KO, or *afadin* KO cells with *ZO-1*/*ZO-2* DKO cells. Data represent the mean ± SD (*n* = 3 each), **p < 0.005, ***p < 0.0005, compared by one-way ANOVA followed by Bonferroni *t*-test. (E) Snapshots from time-lapse imaging of *ZO-1*/*ZO-2* DKO cells expressing EGFP co-cultured with *E-cadherin* KO or *afadin* KO cells. *ZO-1*/*ZO-2* DKO cells were not eliminated. See Fig. 2A for the co-culture with MDCK II cells. (F) Quantification of the numbers of cell deaths (magenta) and the area fraction occupied by *ZO-1*/*ZO-2* DKO cells (green) upon co-culture with *E-cadherin* KO or *afadin* KO cells. The elimination of *ZO-1*/*ZO-2* DKO cells was suppressed by *E-cadherin* or *afadin* KO. See Fig. 2B for the co-culture with MDCK II cells. Scale bars: 20 μm. See also Video S5.

### Hippo signaling in the surrounding cells is required for the elimination of *ZO-1*/*ZO-2* DKO cells

In addition to AJs, Hippo signaling plays an important role in mechanotransduction^37^. In solo culture of parental MDCK II cells, TAZ/YAP, the transcription factor that mediates Hippo signaling, was primarily localized in the cytoplasm (Fig. 6A, F), whereas in solo culture of *ZO-1*/*ZO-2* DKO cells, TAZ/YAP accumulated in the nucleus (Fig. 6B, G), consistent with previous results^38,39^. Interestingly, upon co-culture of parental MDCK II cells with *ZO-1*/*ZO-2* DKO cells, nuclear localization of TAZ/YAP was observed in parental MDCK II cells (Fig. 6C, F), and this was reduced on ROCK or afadin inhibition (Fig. 6D, E–G). These results indicate that the nuclear localization of TAZ/YAP in the surrounding WT cells correlates with their ability to eliminate *ZO-1*/*ZO-2* DKO cells.

**Figure 6.**
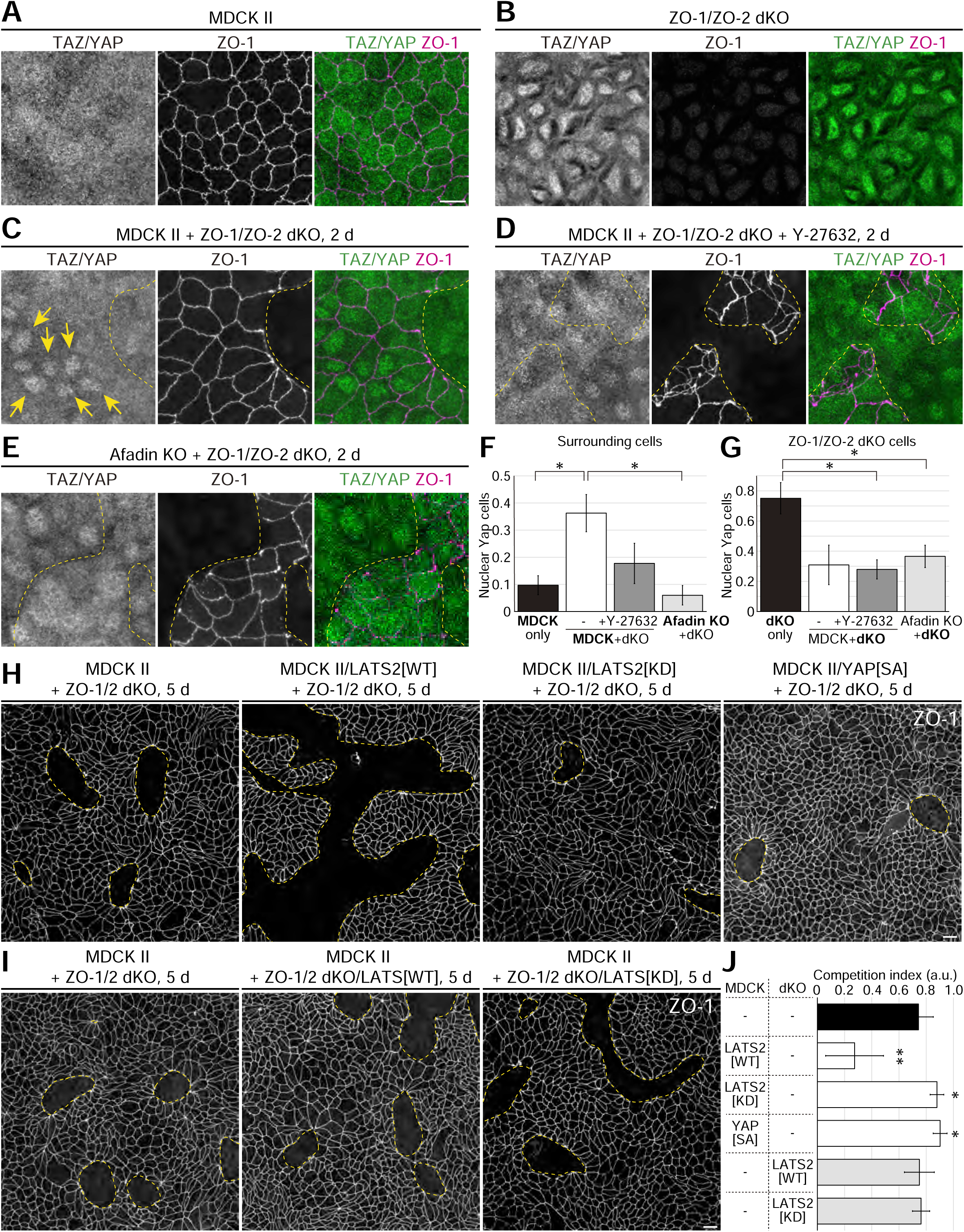
TAZ/YAP nuclear translocation in the surrounding WT cells regulates *ZO-1*/*ZO-2* DKO cell elimination. (A) Cytoplasmic localization of TAZ/YAP in confluent MDCK II cells. (B) Nuclear localization of TAZ/YAP in confluent *ZO-1*/*ZO-2* DKO cells. (C) Localization of TAZ/YAP in the co-culture of MDCK II cells and *ZO-1*/*ZO-2* DKO cells. Nuclear translocation of TAZ/YAP was observed in surrounding MDCK II cells (yellow arrows). (D) Localization of TAZ/YAP in the co-culture of MDCK II cells and *ZO-1*/*ZO-2* DKO cells treated with Y-27632. Nuclear translocation of TAZ/YAP in surrounding MDCK II cells was suppressed by Y-27632 treatment. (E) Localization of TAZ/YAP in the co-culture of *afadin* KO cells and *ZO-1*/*ZO-2* DKO cells. Nuclear translocation of TAZ/YAP was suppressed in surrounding *afadin* KO cells. (F) Quantification of the nuclear localization of TAZ/YAP in the surrounding cells. TAZ/YAP nuclear localization was observed when MDCK II cells were co-cultured with *ZO-1*/*ZO-2* DKO cells, but was suppressed by Y-27632 treatment or *afadin* KO (*n* = 3 each, **p < 0.005, compared by one-way ANOVA with Tukey post-hoc test). (G) Quantification of the nuclear localization of TAZ/YAP in *ZO-1*/*ZO-2* DKO cells. TAZ/YAP nuclear localization was observed in *ZO-1*/*ZO-2* DKO cells, but was suppressed by co-culture with MDCK II cells or *afadin* KO cells, and was not affected by ROCK inhibition (*n* = 3 each, *p < 0.05, compared by one-way ANOVA with Tukey post-hoc test). (H) Hippo signaling in surrounding MDCK II cells regulates the elimination of *ZO-1*/*ZO-2* DKO cells. *ZO-1*/*ZO-2* DKO cell elimination was suppressed by overexpression of LATS2[WT], and promoted by overexpression of LATS2[KD] or YAP[SA]. (I) Hippo signaling in *ZO-1*/*ZO-2* DKO cells is not important for their elimination. Overexpression of LATS[WT] or LATS[KD] did not affect the efficiency of *ZO-1*/*ZO-2* DKO cell elimination. (J) Quantification of the effect of perturbation of Hippo signaling on the efficiency of *ZO-1*/*ZO-2* DKO cell elimination. Data represent the mean ± SD (*n* = 6 each), *p < 0.05, **p < 0.01, compared with MDCK II and *ZO-1*/*ZO-2* DKO co-culture by one-way ANOVA with Tukey post-hoc test. Scale bars: 10 μm (A–E); 20 μm (H, I).

To investigate the role of Hippo signaling in the elimination of *ZO-1*/*ZO-2* DKO cells, we perturbed Hippo signaling. TAZ/YAP nuclear localization is regulated by phosphorylation by an upstream kinase, LATS^40^. When TAZ/YAP is phosphorylated, it localizes to the cytoplasm, and dephosphorylated TAZ/YAP localizes to the nucleus^40^. Overexpression of LATS2[WT] in parental MDCK II cells inhibited the elimination of *ZO-1*/*ZO-2* DKO cells, while overexpression of LATS2[KD] (a kinase-dead form) or YAP[SA] (a phosphorylation-deficient form) promoted the elimination of *ZO-1*/*ZO-2* DKO cells (Fig. 6H, J). By contrast, overexpression of LATS2[WT] or LATS2[KD] in *ZO-1*/*ZO-2* DKO cells did not affect their elimination (Fig. 6I, J). These results suggest that nuclear localization of TAZ/YAP in the surrounding WT cells is required to eliminate *ZO-1*/*ZO-2* DKO cells.

### A ROCK–Hippo–afadin axis regulates the elimination of *ZO-1*/*ZO-2* DKO cells

The above results establish that ROCK, afadin, and Hippo signaling are required to eliminate *ZO-1*/*ZO-2* DKO cells. To examine the relationship between these factors, we performed epistasis analysis. The ROCK inhibitor Y-27632 inhibited the elimination of *ZO-1*/*ZO-2* DKO cells, while overexpression of LATS2[KD] or YAP[SA] in MDCK II cells promoted *ZO-1*/*ZO-2* DKO cell elimination. The elimination of *ZO-1*/*ZO-2* DKO cells by MDCK II cells expressing LATS2[KD] or YAP[SA] was not inhibited by Y-27632 treatment (Fig. 7A, B), demonstrating that Hippo signaling acts downstream of ROCK. By contrast, *ZO-1*/*ZO-2* DKO cell elimination was not restored when LATS2[KD] or YAP[SA] was overexpressed in afadin KO cells (Fig. 7C, D), demonstrating that afadin lies downstream of Hippo signaling. Taken together, these results suggest that a ROCK–Hippo–afadin axis regulates *ZO-1*/*ZO-2* DKO cell elimination (Fig. 7E).

**Figure 7.**
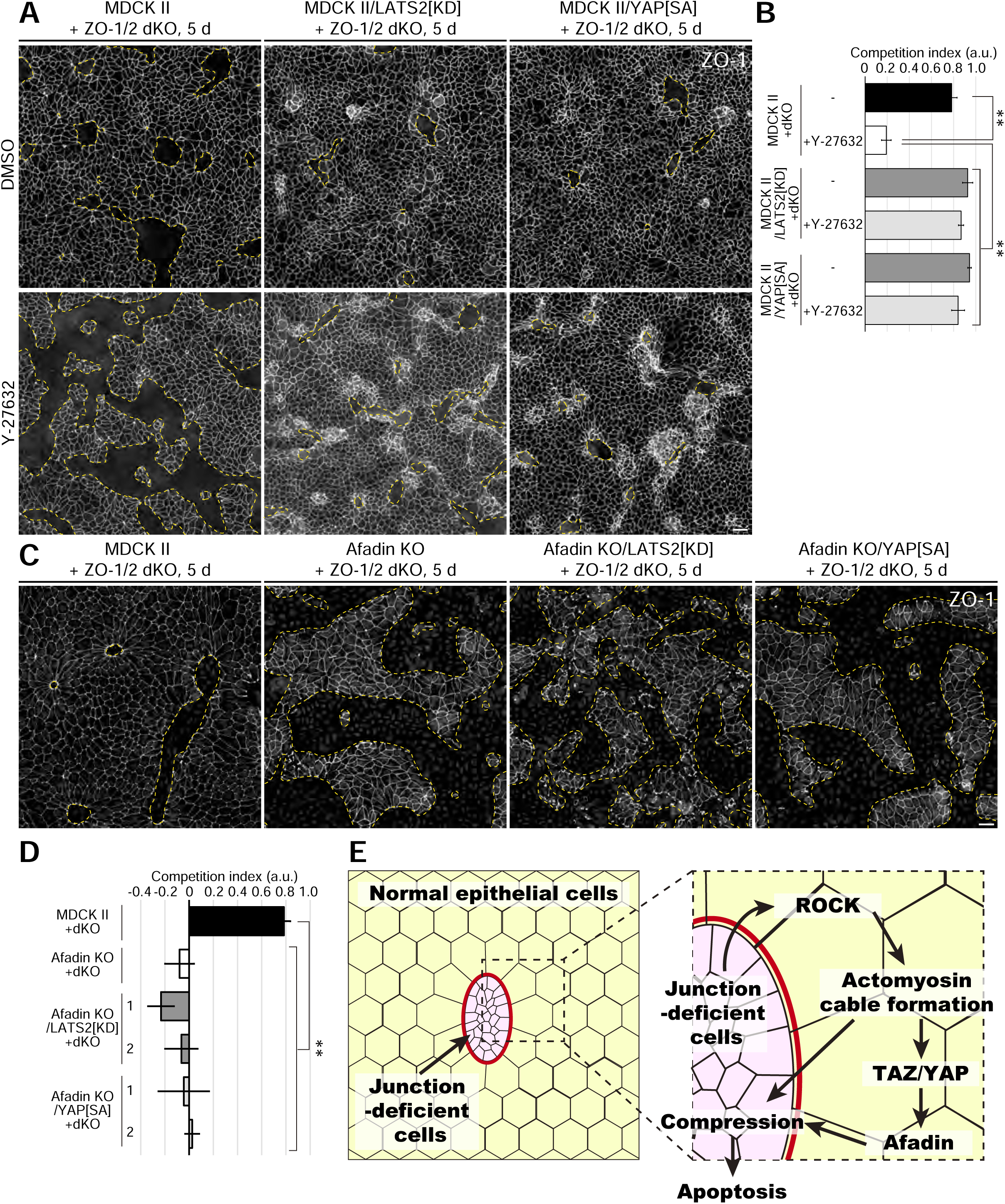
A ROCK–YAP–afadin axis regulates the elimination of *ZO-1*/*ZO-2* DKO cells. (A) Overexpression of LATS2[KD] or YAP[SA] in surrounding MDCK II cells suppressed the effect of Y-27632 treatment. (B) Quantification of the combined effect of Hippo signaling perturbation and Y-27632 treatment on the elimination efficiency of *ZO-1*/*ZO-2* DKO cells. Data represent the mean ± SD (*n* = 5 each), **p < 0.01, compared by one-way ANOVA with Tukey post-hoc test. (C) Overexpression of LATS2[KD] or YAP[SA] in *afadin* KO cells did not restore the ability to eliminate *ZO-1*/*ZO-2* DKO cells. (D) Quantification of the combined effect of perturbation of Hippo signaling and *afadin* KO on the efficiency of *ZO-1*/*ZO-2* DKO cell elimination. Data represent the mean ± SD (*n* = 5 each), **p < 0.01, compared by one-way ANOVA with Tukey post-hoc test. (E) Model of *ZO-1*/*ZO-2* DKO cell elimination. The presence of cell–cell junction-deficient cells (pink) in the epithelial monolayer induces ROCK-dependent formation of supracellular actomyosin cables (red) in the surrounding WT cells at the clonal boundary. This actomyosin cable contracts in a purse-string-like fashion to compress the cell–cell junction-deficient cells, leading to the nuclear translocation of TAZ/YAP and the mechanosensitive accumulation of afadin at the clonal boundaries, promoting the elimination of cell–cell junction-deficient cells by inducing apoptosis. Scale bars: 20 μm.

## Discussion

Epithelia are fundamental in the maintenance of homeostasis of the internal body. The epithelial barrier is challenged by various stresses, including mechanical forces, chemical substances, and infections. However, how epithelial barrier defects are detected and repaired to maintain homeostasis is poorly understood. In this study, we demonstrate that ZO-1/ZO-2-deficient cells are eliminated from epithelia by cell competition, resulting in the restoration of epithelial barrier function. This elimination depends on the interfacial tension generated by neighboring WT cells and is regulated by the ROCK–Hippo–afadin axis. Taken together, these results suggest that cell competition-mediated ZO-1/ZO-2-deficient cell elimination regulates epithelial barrier homeostasis.

### Role of cell competition in epithelial barrier homeostasis

Previous studies have revealed several pathways involved in epithelial barrier homeostasis. For example, local, micrometer-scale breaches of the epithelial barrier are repaired by transient local activation of Rho, which leads to the reassembly of tight junctions through actin polymerization and ROCK-dependent junction contraction^19,41^. In addition, local disruption of the TJ fence function has been shown to induce the cleavage of EpCAM downstream of Rho–ROCK, resulting in release of the reservoir pool of claudins^20,21^. Under inflammatory conditions, junction-inducing peptide (JIP) is produced and is involved in the restoration of TJs via Gα13, which presumably activates Rho signaling^22^. In this study, we have demonstrated that mesoscopic focal epithelial barrier defects are repaired by cell competition. This pathway also involves ROCK signaling, suggesting that Rho signaling plays a central role in regulating epithelial barrier homeostasis by organizing the actomyosin cytoskeleton underlying the apical junctional complex.

Our results suggest that ZO-1/ZO-2-deficient cells are eliminated via mechanical cell competition, where the losing cells are compressed by supracellular actomyosin cables formed by neighboring cells at the clonal interface. In an accompanying paper, Valon et al demonstrate that interfacial tension can drive mechanical competition using Drosophila pupal notum and mathematical modeling^42^. Altogether, these results establish the importance of interfacial tension in mechanical cell competition.

MDCK cells undergo live cell extrusion upon overcrowding^43^. However, in contrast to live cell extrusion, caspase was activated in the ZO-1/ZO-2-deficient cells upon compression, suggesting that TJ-deficient cells could be vulnerable to mechanical stress in a manner similar to *scribble*-KD cells^44^. How mechanical stress triggers apoptosis in ZO-1/ZO-2-deficient cells is an important issue for future investigation. In addition, how the surrounding WT cells recognize ZO-1/ZO-2-deficient cells remains unclear. Because ZO-1/ZO-2-deficient cells show pleiotropic defects, including epithelial barrier dysfunction, epithelial polarity defects, and actomyosin disorganization^6–9^, the functions of ZO-1/ZO-2 should be dissected to determine the trigger for their elimination.

### Molecular mechanisms of epithelial homeostasis

Our results suggest that the elimination of ZO-1/ZO-2-deficient cells involves a purse-string-like constriction of supracellular actomyosin cables formed in the surrounding WT cells at the clone boundary. Previous results have shown that loss of ZO-1 or TJs results in actomyosin cable assembly^9,45–48^, suggesting that focal cell junction defects may be sensed by neighboring cells to trigger actomyosin cable assembly. The supracellular actomyosin cable is similar to that observed during wound healing, apoptotic cell extrusion, or dorsal/thorax closure during *Drosophila* morphogenesis^49–55^. Epithelial wounds, cell extrusion sites, and the leading edge of dorsal/thorax closure are all sites where local mechanical heterogeneities exist within the epithelia. Although the molecular mechanisms underlying supracellular actomyosin cable assembly are still poorly characterized, recent results suggest that cell junction-dependent mechanosensing may play an important role^56^, and focal cell junction defects may be directly sensed by mechanosensors at the cell junctions to regulate epithelial homeostasis. Given that the AJs play central roles in apoptotic cell extrusion and the elimination of ZO-1/ZO-2-deficient cells, it will be interesting to see whether common molecular machineries underlie these processes.

Our results indicate that the mechanosensitive recruitment of afadin to the clone boundary is important for the elimination of ZO-1/ZO-2-deficient cells. Afadin has been suggested to act as a mechanosensor^57,58^. Given that the localization of afadin is regulated by various domains in a complex manner^59–65^, the functional domains involved in the accumulation of afadin at the clonal boundary in response to tension should be investigated in the future. Afadin plays a key role in anchoring the actomyosin cable to the AJs^66,67^, implying that afadin may promote the elimination of ZO-1/ZO-2-deficient cells by reinforcing the purse-string-like contraction, by coupling the supracellular actomyosin cables to the cell–cell junctions.

Our results suggest that TAZ/YAP translocates to the nucleus in response to focal cell junction defects and promotes the elimination of TJ-deficient cells. Although Hippo signaling in neighboring cells was required for the elimination of ZO-1/ZO-2-deficient cells, Hippo signaling perturbation in the losing cells did not alter the phenotype. These results suggest that ZO-1/ZO-2-deficient cell elimination is not caused by Hippo-dependent cell competition, i.e., the Hippo signaling levels in the winning and losing cells are not compared^32,68^. Our results are in line with the hypothesis that TAZ/YAP acts as an epithelial damage sensor^69^. Indeed, increased TAZ/YAP nuclear translocation occurs at the proximity of wound edges and in response to tissue injury^70^. Although it is unclear how TAZ/YAP responds to focal epithelial barrier defects, ZO proteins affect the mechanics of epithelial cells^71^ and Hippo signaling is involved in mechanotransduction^37^, suggesting that mechanical force could be involved in this process.

### Epithelial barrier homeostasis in tissue physiology

Epithelial barrier function is compromised in various inflammatory diseases and cancers^5,72^. Interestingly, previous results have shown that epithelial cells infected with *Listeria monocytogenes* or *Rickettsia parkeri* are collectively extruded from epithelial monolayers, and cell–cell adhesion and the cytoskeleton are involved in this process^73^. In addition, cells transformed by Ras[V12] or v-Src exhibit disorganized epithelial structures and are extruded from epithelial monolayers by cell competition^26,74–78^, and inflammation affects epithelial barrier function and is frequently accompanied by epithelial shedding^79–81^. In summary, these results suggest that removal of cell–cell junction-deficient cells by cell competition may be a common mechanism to maintain epithelial homeostasis, and it will be of interest to investigate whether common molecular mechanisms underlie these processes. Identifying the molecular markers of cell competition that regulate the elimination of cell–cell junction-deficient cells will greatly facilitate our understanding of the roles of cell competition in epithelial homeostasis in physiological contexts.

In conclusion, we have demonstrated that ZO-1/ZO-2-deficient cells are eliminated from epithelia by cell competition. Further investigation of the molecular mechanisms underlying the recognition of ZO-1/ZO-2-deficient cells, and how mechanical stress induces apoptosis in ZO-1/ZO-2-deficient cells, should deepen our understanding of how homeostasis of the epithelial barrier is maintained.

## Acknowledgments

We thank Akira Nagafuchi, Masayuki Murata, Ernst Reichmann, Clark Distelhorst, Michael Davidson, Hiroshi Hosoya, Hiroshi Sasaki, Manabu Kawada, and the Developmental Studies Hybridoma Bank for kindly providing reagents; Mika Watanabe and Yuichiro Kano for technical assistance; the Bioimaging Facility of NIBB and Misako Saida for their support in laser ablation experiments; Kaoru Sugimura for advice on laser ablation experiments and image analyses; Toshiro Moroishi for advice on TAZ/YAP immunostaining; and all members of the Furuse laboratory and Otani laboratory for discussions and comments. We also thank Catherine Perfect from Edanz for editing a draft of this manuscript.

This work was supported by a JSPS Grants-in-Aid for Challenging Exploratory Research (16K15226, to MF), JSPS Grants-in-Aid for Scientific Research (B) (26291043, 18H02440, 21H02523, to MF), JSPS Grants-in-Aid for Young Scientists (B) (16K18544, to TO), JSPS Grants-in-Aid for Scientific Research (C) (18K06234, TO), MEXT/JSPS Grants-in-Aid for Scientific Research on Innovative Areas (17H05627, to TO), MEXT/JSPS Grants-in-Aid for Transformative Research Areas (21H05286, to TO), JST PRESTO (JPMJPR204, to TO), MEXT/JSPS Grants-in-Aid for Scientific Research on Innovative Areas – Platforms for Advanced Technologies and Research Resources “Advanced Bioimaging Support” (JP16H06280, JP22H04926), Molecular Profiling Committee, MEXT/JSPS Grants-in-Aid for Scientific Research on Innovative Areas “Advanced Animal Model Support (AdAMS) (JP16H06276), the Inamori Foundation (to TO), the Japan Spina Bifida & Hydrocephalus Research Foundation (to TO), and the Takeda Science Foundation (to MF, TO).

## Author contributions

Conceptualization, Funding acquisition, Supervision, Project administration: T.O. and M.F.; Resources: T.O., T.P.N., N.K., T.F., and M.F.; Investigation, Formal analysis, Visualization: T.O., T.P.N., N.K.; Writing – original draft: T.O. and M.F.; Writing – review & editing: all authors.

## Supplemental information titles and legends

**Figure S1.**
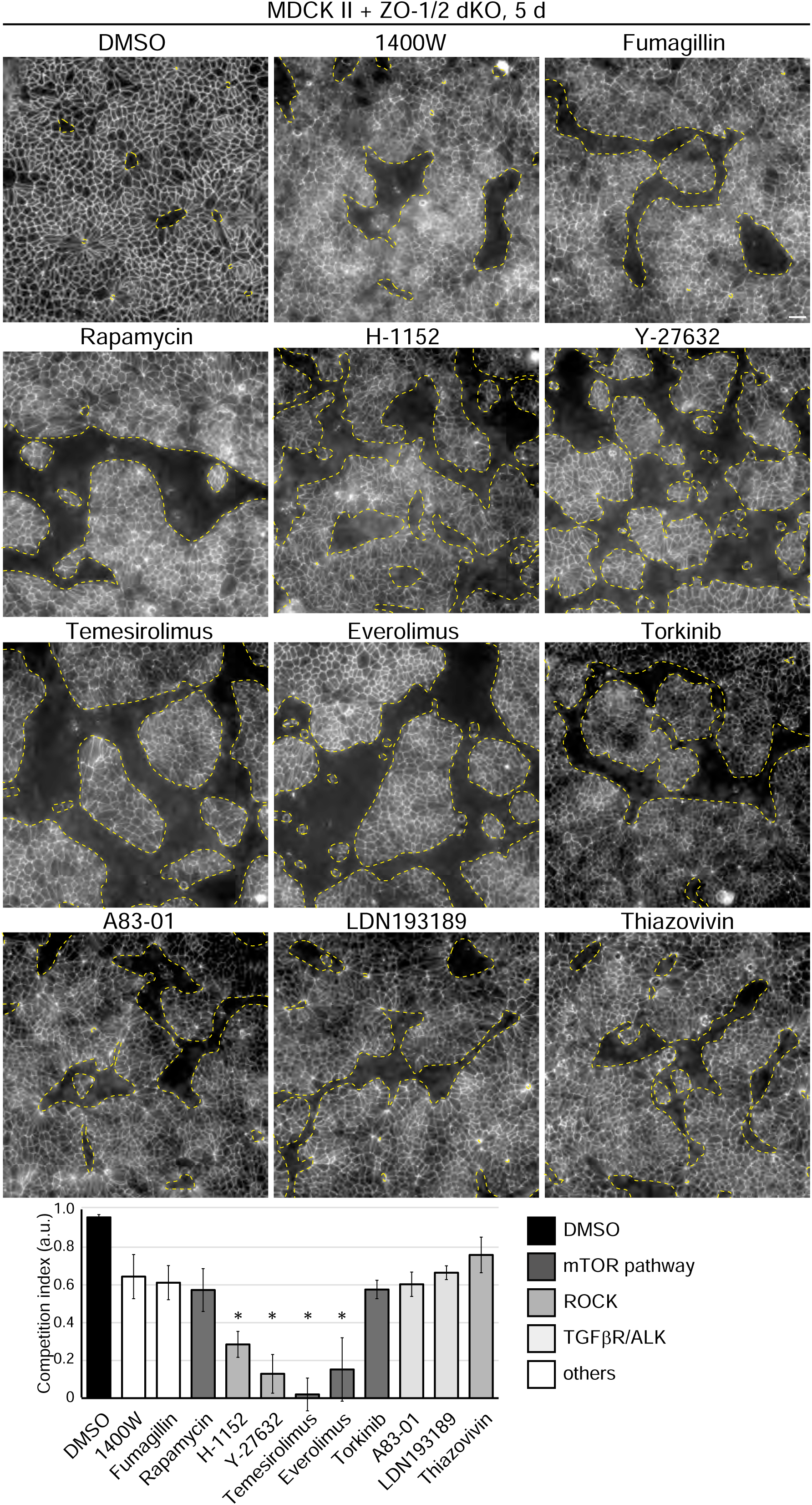
Representative results of the chemical compound library screening. Representative images of anti-ZO-1 staining of the co-culture of MDCK II cells and *ZO-1*/*ZO-2* DKO cells treated with various chemical compounds. The concentrations used were all 10 μM, except for leptomycin B, which was used at 0.2 mM. Data represent the mean ± SD (*n* = 3 each), *p < 0.05, compared by one-way ANOVA with Tukey post-hoc test. Scale bar: 20 μm.

**Figure S2.**
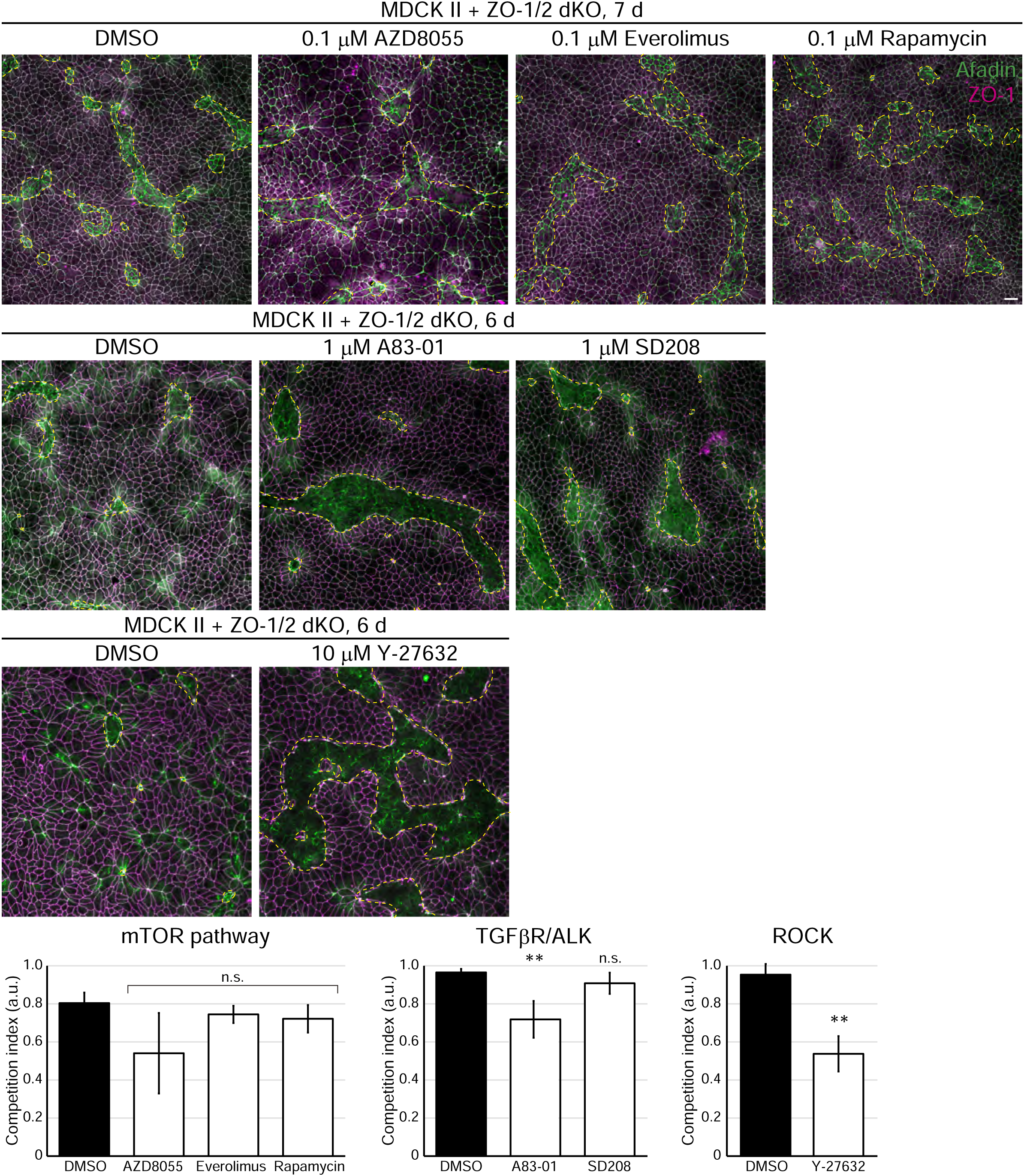
Y-27632 inhibits the elimination of *ZO-1*/*ZO-2* DKO cells. Representative images of anti-ZO-1 staining of the co-culture of MDCK II cells and *ZO-1*/*ZO-2* DKO cells treated with inhibitors of the mTOR pathway (AZD8055, everolimus, rapamycin), the TGFβR/ALK pathway (A83-01, SD208), or the ROCK pathway (Y-27632). The data represent the mean ± SD (*n* = 3 each), **p < 0.01, compared by one-way ANOVA with Tukey post-hoc test. Scale bar: 20 μm.

**Figure S3.**
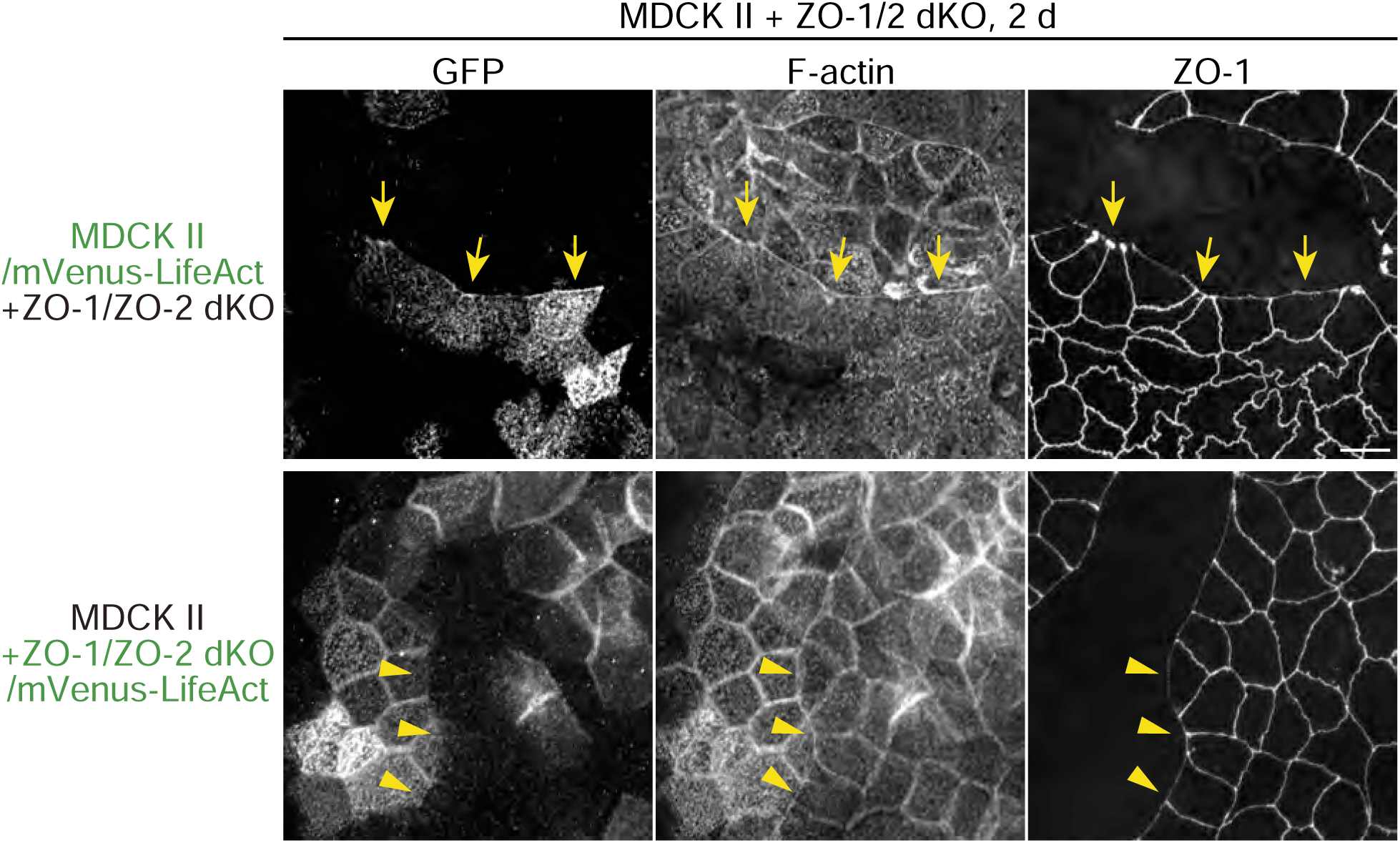
Actin cables form at the clone boundary in surrounding WT cells. The supracellular actomyosin cable is formed in the surrounding WT cells. The supracellular actomyosin cables were labeled by mVenus-LifeAct when expressed in MDCK II cells (yellow arrows) but not when expressed in *ZO-1*/*ZO-2* DKO cells (yellow arrowheads), suggesting that the actomyosin cables formed in the surrounding WT cells. Scale bar: 10 μm.

**Figure S4.**
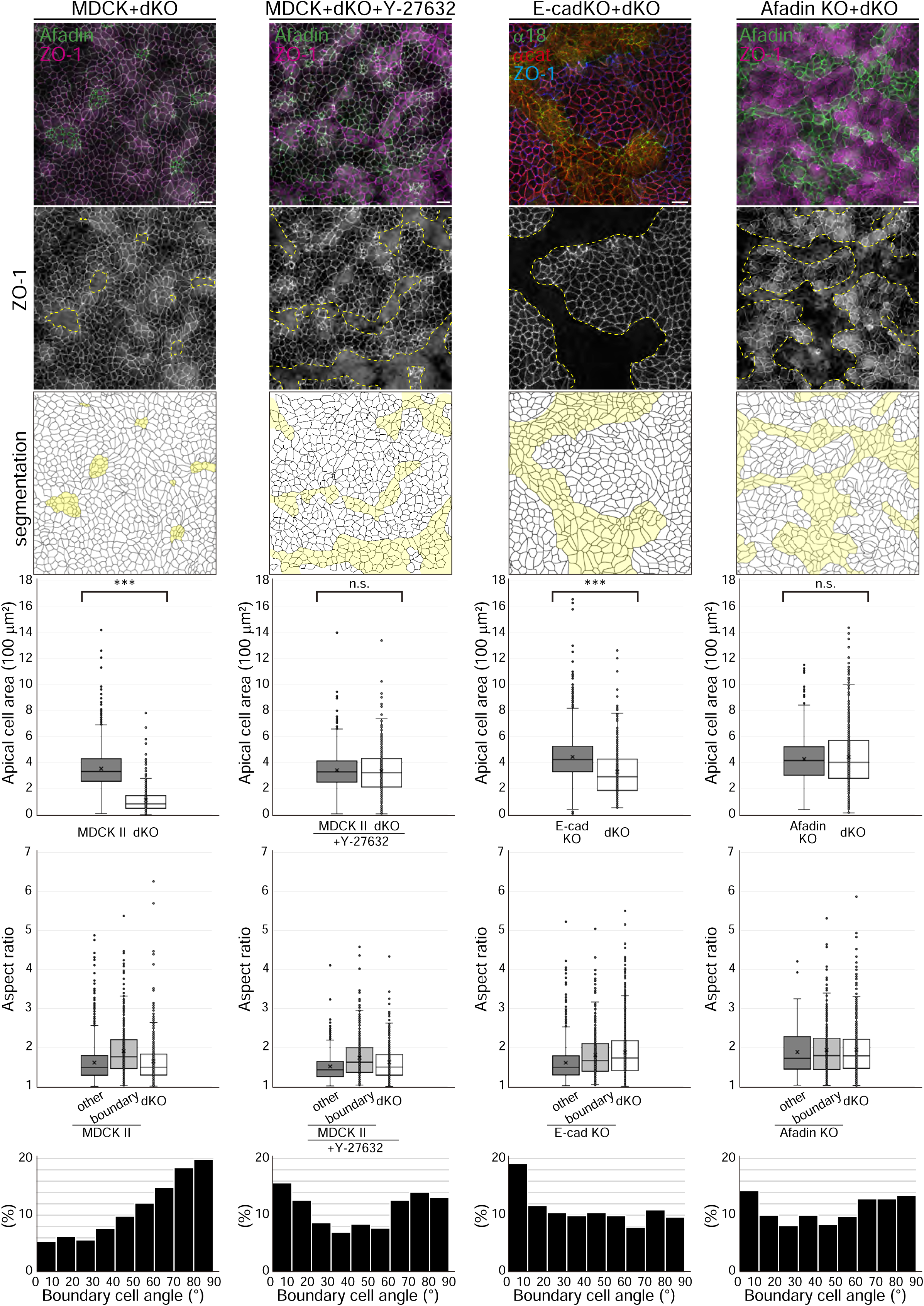
*ZO-1*/*ZO-2* DKO cells are compressed. Cell outlines were segmented using cell–cell junction marker immunostaining of MDCK II and *ZO-1*/*ZO-2* DKO cell co-culture, MDCK II and *ZO-1*/*ZO-2* DKO cell co-culture treated with Y-27632, *E-cadherin* KO and *ZO-1*/*ZO-2* DKO cell co-culture, and *afadin* KO and *ZO-1*/*ZO-2* DKO cell co-culture. The apical cell area of the surrounding cells and the *ZO-1*/*ZO-2* DKO cells were measured. *ZO-1*/*ZO-2* DKO cells were significantly compressed when co-cultured with MDCK II cells. The compression of *ZO-1*/*ZO-2* DKO cells was completely suppressed by Y-27632 treatment or co-culture with *afadin* KO cells, and was partially suppressed by co-culture with *E-cadherin* KO cells (*n* = 2553 and 530 for MDCK II and *ZO-1*/*ZO-2* DKO cells, respectively; *n* = 803 and 532 for MDCK II and Y-27632-treated *ZO-1*/*ZO-2* DKO cells, respectively; *n* = 1620 and 1268 for *E-cadherin* KO and *ZO-1*/*ZO-2* DKO cells, respectively; *n* = 617 and 511 for *afadin* KO and *ZO-1*/*ZO-2* DKO cells, respectively), ***p < 0.0005, compared by *t*-test. The aspect ratio of each cell was calculated by ellipsoid fitting and measurement of the ratio between the major and minor axis. MDCK II cells facing the clonal boundary adopted elongated morphologies, while this cell elongation was suppressed by knockout of E-cadherin or afadin (*n* = 1856, 678, and 530 for MDCK II cells (other), MDCK II cells (boundary), and *ZO-1*/*ZO-2* DKO cells, respectively; *n* = 396, 407, and 532 for MDCK II cells (other), MDCK II cells (boundary), and Y-27632-treated *ZO-1*/*ZO-2* DKO cells, respectively; *n* = 1228, 392, and 1268 for *E-cadherin* KO cells (other), *E-cadherin* KO cells (boundary), and *ZO-1*/*ZO-2* DKO cells, respectively; *n* = 191, 426, and 511 for *afadin* KO cells (other), *afadin*-KO cells (boundary), and *ZO-1*/*ZO-2* DKO cells, respectively). The axis of the clone boundary cells relative to the clonal boundary was measured by measuring the angle between the major axis of the cell and the clone boundary. MDCK II cells were oriented perpendicular to the clonal boundary, whereas no such directionality was observed on Y-27632 treatment or knockout of E-cadherin or afadin. Scale bars: 20 μm.

**Video S1. *ZO-1*/*ZO-2* DKO cells are eliminated upon co-culture with MDCK II cells.**

*ZO-1*/*ZO-2* DKO cells expressing EGFP were co-cultured with MDCK II cells for 5 d. Videos were taken at 10 min intervals with z-stacking with 2 μm intervals for 20 μm. The movie shows maximum intensity projection images. EGFP-positive *ZO-1*/*ZO-2* DKO cells were progressively eliminated by apoptosis.

**Video S2. Purse-string-like contraction of the supracellular actomyosin cables surrounding *ZO-1*/*ZO-2* DKO cells.**

MDCK II cells expressing MRLC2-EGFP were co-cultured with *ZO-1*/*ZO-2* DKO cells for 5 d. Videos were taken at 10 min intervals with z-stacking with 2 μm intervals for 20 μm. The movie shows maximum intensity projection images. EGFP-labeled actomyosin cables constricted in a purse-string-like manner.

**Video S3. Clonal boundaries are under high tension.**

MDCK II cells expressing mZO-1-EGFP were co-cultured with *ZO-1*/*ZO-2* DKO cells and the cell boundaries between MDCK II cells (left) or the clonal boundaries (right) were laser ablated. Videos were taken at 1 s intervals. The movie is a single focal plane. The clonal boundaries showed greater initial recoil velocity, demonstrating that the clonal boundaries are under higher tension.

**Video S4. *ZO-1*/*ZO-2* DKO cells must be surrounded by MDCK II cells to be eliminated.**

MDCK II cells expressing EGFP were co-cultured with *ZO-1*/*ZO-2* DKO cells expressing mCherry for 5 d. Videos were taken at 10 min intervals with z-stacking with 2 μm intervals for 20 μm. The movie shows maximum intensity projection images. mCherry-positive *ZO-1*/*ZO-2* DKO cells were progressively eliminated by apoptosis when they were surrounded by MDCK II cells (left: co-culture ratio 9:1), but did not undergo cell death when the clonal topology was inverted (right: co-culture ratio 1:9).

**Video S5. AJs in MDCK II cells are required for the elimination of *ZO-1*/*ZO-2* DKO cells.**

*ZO-1*/*ZO-2* DKO cells expressing EGFP were co-cultured with MDCK II cells (left), *E-cadherin* KO cells (center), or *afadin* KO cells (right) for 5 d. Videos were taken at 10 min intervals with z-stacking with 2 μm intervals for 20 μm. The movie shows maximum intensity projection images. EGFP-positive *ZO-1*/*ZO-2* DKO cells were progressively eliminated when co-cultured with MDCK II cells, but the elimination was greatly reduced upon co-culture with *E-cadherin* KO cells or *afadin* KO cells.

## STAR* METHODS

### Key Resources Table

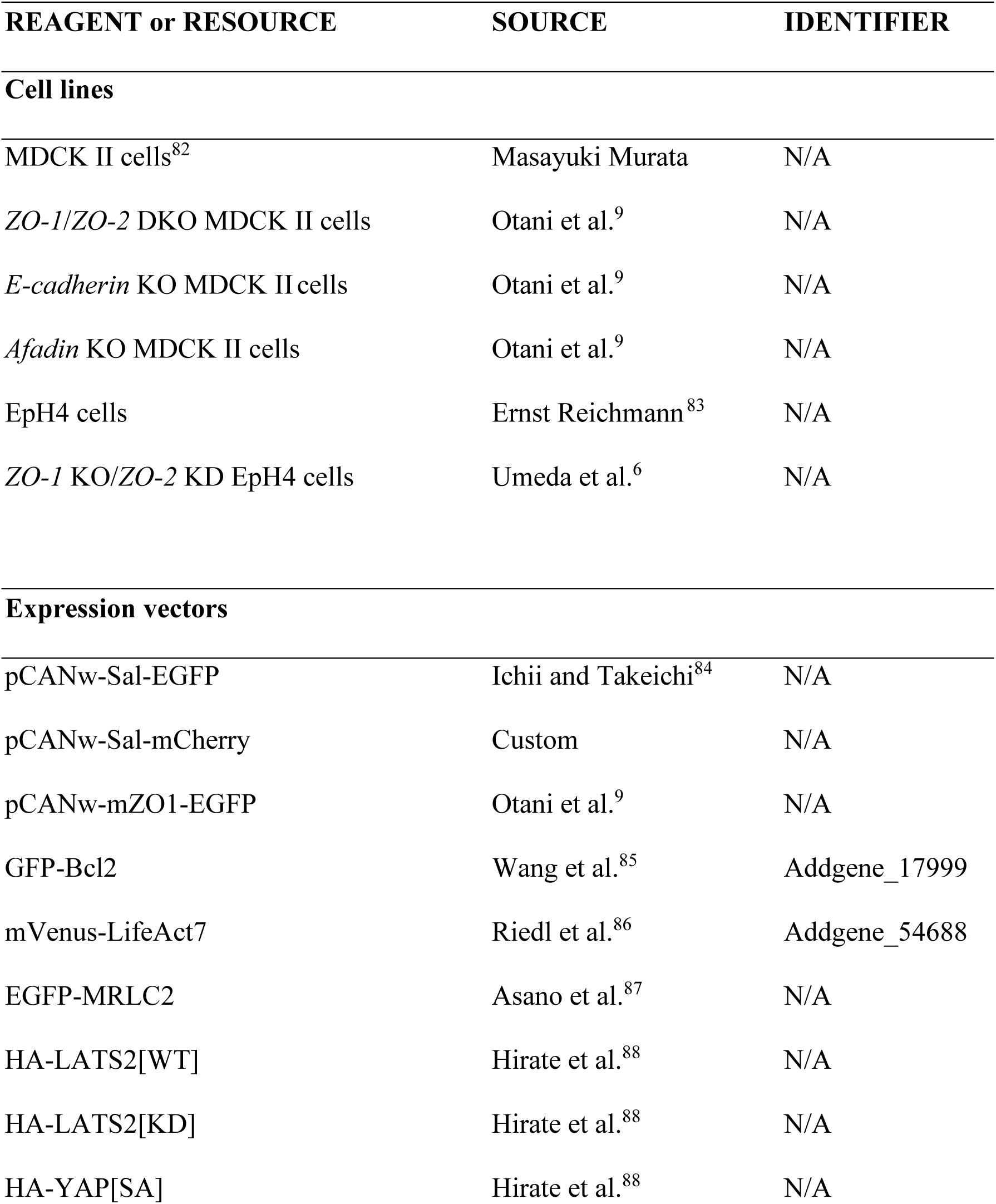
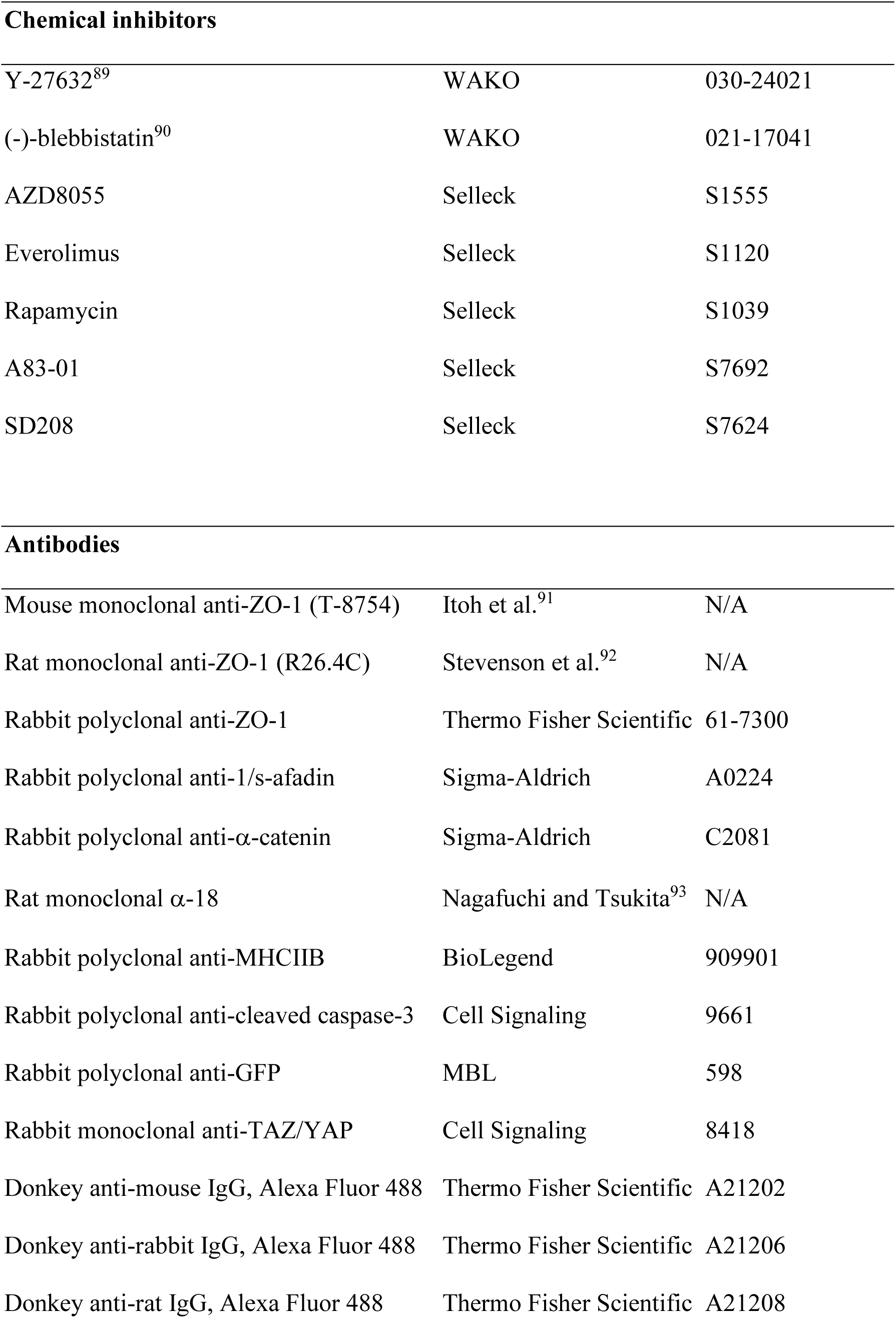
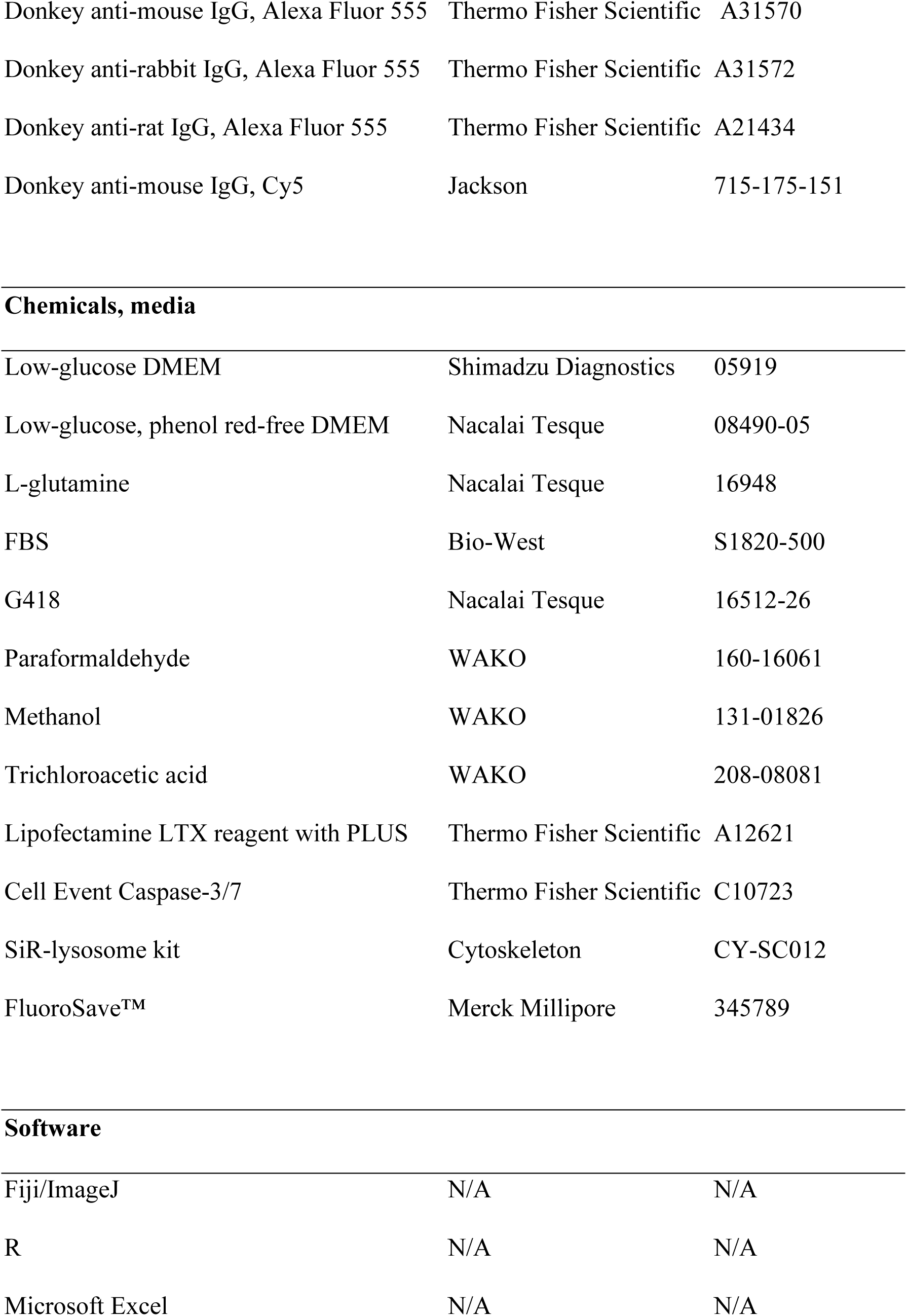

| REAGENT or RESOURCE | SOURCE | IDENTIFIER |
| --- | --- | --- |
| **Cell lines** |  |  |
| MDCK II cells<sup>82</sup> | Masayuki Murata | N/A |
| *ZO-1/ZO-2* DKO MDCK II cells | Otani et al.<sup>9</sup> | N/A |
| *E-cadherin* KO MDCK II cells | Otani et al.<sup>9</sup> | N/A |
| *Afadin* KO MDCK II cells | Otani et al.<sup>9</sup> | N/A |
| EpH4 cells | Ernst Reichmann<sup>83</sup> | N/A |
| *ZO-1* KO/*ZO-2* KD EpH4 cells | Umeda et al.<sup>6</sup> | N/A |
| **Expression vectors** |  |  |
| pCANw-Sal-EGFP | Ichii and Takeichi<sup>84</sup> | N/A |
| pCANw-Sal-mCherry | Custom | N/A |
| pCANw-mZO1-EGFP | Otani et al.<sup>9</sup> | N/A |
| GFP-Bcl2 | Wang et al.<sup>85</sup> | Addgene_17999 |
| mVenus-LifeAct7 | Riedl et al.<sup>86</sup> | Addgene_54688 |
| EGFP-MRLC2 | Asano et al.<sup>87</sup> | N/A |
| HA-LATS2[WT] | Hirate et al.<sup>88</sup> | N/A |
| HA-LATS2[KD] | Hirate et al.<sup>88</sup> | N/A |
| HA-YAP[SA] | Hirate et al.<sup>88</sup> | N/A |

|  |  |  |  |
| --- | --- | --- | --- |
| 691 | **Chemical inhibitors** |  |  |
| 692 | Y-27632<sup>89</sup> | WAKO | 030-24021 |
| 693 | (-)-blebbistatin<sup>90</sup> | WAKO | 021-17041 |
| 694 | AZD8055 | Selleck | S1555 |
| 695 | Everolimus | Selleck | S1120 |
| 696 | Rapamycin | Selleck | S1039 |
| 697 | A83-01 | Selleck | S7692 |
| 698 | SD208 | Selleck | S7624 |
| 699 |  |  |  |
| 700 | **Antibodies** |  |  |
| 701 | Mouse monoclonal anti-ZO-1 (T-8754) | Itoh et al.<sup>91</sup> | N/A |
| 702 | Rat monoclonal anti-ZO-1 (R26.4C) | Stevenson et al.<sup>92</sup> | N/A |
| 703 | Rabbit polyclonal anti-ZO-1 | Thermo Fisher Scientific | 61-7300 |
| 704 | Rabbit polyclonal anti-1/s-afadin | Sigma-Aldrich | A0224 |
| 705 | Rabbit polyclonal anti-α-catenin | Sigma-Aldrich | C2081 |
| 706 | Rat monoclonal α-18 | Nagafuchi and Tsukita<sup>93</sup> | N/A |
| 707 | Rabbit polyclonal anti-MHCIIB | BioLegend | 909901 |
| 708 | Rabbit polyclonal anti-cleaved caspase-3 | Cell Signaling | 9661 |
| 709 | Rabbit polyclonal anti-GFP | MBL | 598 |
| 710 | Rabbit monoclonal anti-TAZ/YAP | Cell Signaling | 8418 |
| 711 | Donkey anti-mouse IgG, Alexa Fluor 488 | Thermo Fisher Scientific | A21202 |
| 712 | Donkey anti-rabbit IgG, Alexa Fluor 488 | Thermo Fisher Scientific | A21206 |
| 713 | Donkey anti-rat IgG, Alexa Fluor 488 | Thermo Fisher Scientific | A21208 |
| 714 | Donkey anti-mouse IgG, Alexa Fluor 555 | Thermo Fisher Scientific | A31570 |
| 715 | Donkey anti-rabbit IgG, Alexa Fluor 555 | Thermo Fisher Scientific | A31572 |
| 716 | Donkey anti-rat IgG, Alexa Fluor 555 | Thermo Fisher Scientific | A21434 |
| 717 | Donkey anti-mouse IgG, Cy5 | Jackson | 715-175-151 |

**Chemicals, media**
|  |  |  |  |
| --- | --- | --- | --- |
| 720 | Low-glucose DMEM | Shimadzu Diagnostics | 05919 |
| 721 | Low-glucose, phenol red-free DMEM | Nacalai Tesque | 08490-05 |
| 722 | L-glutamine | Nacalai Tesque | 16948 |
| 723 | FBS | Bio-West | S1820-500 |
| 724 | G418 | Nacalai Tesque | 16512-26 |
| 725 | Paraformaldehyde | WAKO | 160-16061 |
| 726 | Methanol | WAKO | 131-01826 |
| 727 | Trichloroacetic acid | WAKO | 208-08081 |
| 728 | Lipofectamine LTX reagent with PLUS | Thermo Fisher Scientific | A12621 |
| 729 | Cell Event Caspase-3/7 | Thermo Fisher Scientific | C10723 |
| 730 | SiR-lysosome kit | Cytoskeleton | CY-SC012 |
| 731 | FluoroSave™ | Merck Millipore | 345789 |

**Software**
|  |  |  |  |
| --- | --- | --- | --- |
| 734 | Fiji/ImageJ | N/A | N/A |
| 735 | R | N/A | N/A |
| 736 | Microsoft Excel | N/A | N/A |

### Resource Availability

#### Lead contact

Requests for further information and resources should be directed to and will be fulfilled by the lead contact, Tetsuhisa Otani.

#### Materials availability

All unique/stable reagents generated in this study are available from the lead contact on request.

#### Data and code availability

All data reported in this paper will be shared by the lead contact on request. This paper does not report any original code.

#### Experimental model and study participant details

A canine kidney-derived epithelial cell line, MDCK II cells^82^, were generously provided by Masayuki Murata (Institute of Science, Tokyo). *ZO-1*/*ZO-2* DKO MDCK II cells, *E-cadherin* KO MDCK II cells, and *afadin* KO MDCK II cells were as described previously^9^. A mouse mammary tumor-derived epithelial cell line, EpH4 cells, were a gift from Ernst Reichmann (University of Zurich)^83^. *ZO-1* KO/*ZO-2* KD EpH4 cells were as described previously^6^. All cells were maintained in low-glucose DMEM (Shimadzu Diagnostics; #05919) supplemented with 2 mM L-glutamine (Nacalai Tesque; # 16948) and 10% FBS (Bio-West; #S1820-500) at 37°C under conditions of 5% CO2. The medium was changed every 1–2 d.

### Method details

#### Cell culture and drug treatments

Co-culture was performed by mixing the cells at a 1:1 ratio upon plating. The ROCK inhibitor Y-27632 (Wako; # 030-24021)^90^ was used at 10 μM, and the myosin II inhibitor (−)-blebbistatin (Wako; #021-17041)^91^ was used at 50 μM. The drugs were added at the beginning of co-culture and the cells were incubated in the presence of the drug until fixation. The SCADS inhibitor kit was provided by Manabu Kawada. The following drugs were used to reanalyze the pathways that gave positive hits in the first screening: AZD8055 (Selleck; #S1555); everolimus (Selleck, #S1120); rapamycin (Selleck, #S1039); A83-01 (Selleck; #S7692); and SD208 (Selleck, #S7624).

#### Molecular biology

The following mammalian expression vectors were used in this study: pCANw-Sal-EGFP^84^; pCANw-Sal-mCherry, wherein the EGFP gene of pCANw-Sal-EGFP was replaced with an mCherry gene; pCANw-mZO1-EGFP^8^; GFP-BCL2, from Clark Distelhorst (Addgene plasmid #17999)^85^; mVenus-LifeAct7, provided by Michael Davidson (Addgene plasmid #54688)^86^; EGFP-MRLC2, a gift from Hiroshi Hosoya (Kanagawa University)^87^; and HA-LATS2[WT], HA-LATS2[KD], and HA-YAP[SA], generously shared by Hiroshi Sasaki (Osaka University)^88^.

Stable transfectants were established by transfection of the expression vectors using Lipofectamine® LTX reagent with PLUS™ reagent (Thermo Fisher Scientific; #A12621) according to the manufacturer’s instructions, and selected by 400 μg/ml G418 (Nacalai Tesque; #16512-26). Multiple surviving clones were isolated, and the expression of each transgene was confirmed by immunofluorescence or western blotting. Multiple clones were isolated for each construct, and all clones showed consistent phenotypes.

#### Antibodies

The following primary antibodies were used for immunofluorescence analyses: mouse monoclonal anti-ZO-1 (clone T-8754)^91^; rat monoclonal anti-ZO-1 (R26.4C) obtained from Developmental Studies Hybridoma Bank^92^; rabbit polyclonal anti-ZO-1 (Thermo Fisher Scientific; #61-7300); rabbit polyclonal anti-1/s-afadin (Sigma-Aldrich; #A0224); rabbit polyclonal anti-α-catenin (Sigma-Aldrich; #C2081); rat monoclonal α-18, which recognizes the tension-sensitive epitope of α-catenin (kindly provided by Akira Nagafuchi)^10,93^; rabbit polyclonal anti-MHCIIB (BioLegend; #909901); rabbit polyclonal cleaved caspase-3 (Cell Signaling Technology; #9661); rabbit polyclonal anti-GFP (MBL; #598); and rabbit monoclonal anti-TAZ/YAP (Cell Signaling Technology; #8418).

The following secondary antibodies were used to detect the primary antibodies: Alexa Fluor 488-conjugated donkey anti-mouse IgG (Thermo Fisher Scientific; #A21202); Alexa Fluor 488-conjugated donkey anti-rabbit IgG (Thermo Fisher Scientific; #A21206); Alexa Fluor 488-conjugated donkey anti-rat IgG (Thermo Fisher Scientific; #A21208), Alexa Fluor 555-conjugated donkey anti-mouse IgG (Thermo Fisher Scientific; #A31570); Alexa Fluor 555-conjugated donkey anti-rabbit IgG (Thermo Fisher Scientific; #A31572); Alexa Fluor 555-conjugated goat anti-rat IgG (Thermo Fisher Scientific; #A21434); and Cy5-conjugated donkey anti-mouse IgG (#715-175-151; Jackson ImmunoResearch Laboratories). Caspase 3/7 activity was detected by CellEvent Caspase-3/7 (Thermo Fisher Scientific; #C10723), and lysosomes were visualized by an SiR-lysosome kit (Cytoskeleton, Inc.; #CY-SC012).

#### Immunofluorescence

For immunofluorescence, 1 × 10^5^ cells were cultured on Transwell® polycarbonate filters (0.4 µm pore size; Corning; #3401) for 1–7 d. Cells were fixed with 1% PFA (RT, 5 min) or 100% methanol (−20°C, 10 min). Specifically, 1% PFA fixation was used for α-18, anti-cleaved caspase-3, anti-GFP, and anti-TAZ/YAP, and methanol fixation was used for the others. After fixation, the cells were washed three times with PBS, permeabilized with 0.1% Triton X-100 in PBS (RT, 15 min), rinsed once with PBS, and blocked with 10% FBS (RT, 30 min). For cells cultured on Transwell filters, the filters were excised using scalpels before blocking. All cells were incubated with the primary antibodies diluted in blocking solution (RT, 1 h), washed three times with PBS, and incubated with secondary antibodies diluted in blocking solution (RT, 1 h). Finally, the cells were washed three times with PBS and mounted using FluoroSave™ Reagent (Merck Millipore; #345789).

#### Confocal imaging

Confocal imaging was performed by a Leica TCS-SPE confocal laser scanning microscope system mounted on a DMI 4000B inverted microscope equipped with HCX PL FLUOTAR 40× (NA0.75), HCX PL APO 63× (NA1.40), or HCX FL APO 100×(NA1.40) objective lenses, diode lasers (488/532/635 nm), and the LAS AF software (all from Leica Microsystems), or by an AX R confocal laser scanning microscope system mounted on an Eclipse Ti2 inverted microscope equipped with a CFI PLAN Apochromat lambda D 60× oil (NA 1.42) objective, diode lasers (488/561/640 nm), and the NIS-Elements C Imaging software (all from Nikon Solutions).

#### Time-lapse imaging

For time-lapse imaging, 2.5 × 10^6^ cells were plated in glass-based dishes (Iwaki; #3910-035) in low glucose, phenol red-free DMEM (Nacalai Tesque; #08490-05) supplemented with 10% FBS and penicillin-streptomycin-glutamine (Gibco; #10378-016). Time-lapse imaging was performed immediately after plating using a CellVoyager CV1000 spinning disc confocal imaging system (Yokogawa) with a UPLSApo20× dry (NA 0.75) objective lens (Evident) and diode lasers (488/561 nm). The cells were imaged at 10 min intervals for 5 d with z-sectioning at 2 µm step size with 20 µm range. Movie acquisition was performed using CV1000 software (Yokogawa).

#### Laser ablation

Laser ablation was performed using a spinning disc confocal microscope, CSU-X1 (Yokogawa), mounted on an IX81 inverted microscope (Evident) with an UPLSAPO 60× W/NA1.20 water immersion objective (Evident) and a stage-top incubator (Tokai Hit; # INUBSF-FI-RK). MDCK II cells expressing ZO-1-GFP co-cultured with *ZO-1*/*ZO-2* DKO cells were imaged, and laser ablation was performed by an N2 Micropoint laser (16 Hz, 365 nm, 3 μW; Photonic Instruments). Images were acquired by an EMCCD camera (Andor; #DU-897) at 400 ms intervals, starting from 4 s before the laser ablation to 30 s after the laser ablation. Laser ablation took 1 s, and is not included in the movie.

#### TER measurements

Cells were cultured on Transwell® polycarbonate filters (0.4-μm pore size; #3413; Corning) for 1–7 d. Cells were equilibrated to RT before measurements, and the electric resistance between the apical and basolateral sides was measured by a Millicell ERS-2 Volt-Ohm Meter (Merck Millipore). Blank measurements were performed on Transwell® filters filled with medium without cells. After subtraction of the mean blank values, the electric resistance was multiplied by the area of the Transwell® filter to yield the unit area resistance (Ω × cm^2^).

#### Quantification and statistical analysis Quantitative image analyses

Image analyses and processing were performed using Fiji/ImageJ 1.53t software (National Institutes of Health). Median filter, gaussian filter, or brightness and contrast adjustments were applied for qualitative presentation of images when necessary, without any nonlinear adjustments.

The area occupied by *ZO-1*/*ZO-2* DKO cells was measured from the time-lapse movies by binarizing the GFP signals that labeled the *ZO-1*/*ZO-2* DKO cells and measuring the area of any GFP-positive regions using Fiji/ImageJ. The numbers of cell deaths were manually counted from the time-lapse movies.

Competition indices were calculated by the following formula: (A^MDCK^ ^II^ – A^KO^) / (A^MDCK^ ^II^ + A^KO^), where A^MDCK^ ^II^ is the area occupied by surrounding cells, while A^KO^ is the area occupied by *ZO-1*/*ZO-2* DKO cells. The area occupied by the surrounding cells and *ZO-1*/*ZO-2* DKO cells were determined on the basis of the ZO-1 immunostaining images. The competition index will be +1.0 when the surrounding cells completely eliminate the *ZO-1*/*ZO-2* DKO cells, 0 when the surrounding cells and the *ZO-1*/*ZO-2* DKO cells are neutral to each other, and −1.0 when the *ZO-1*/*ZO-2* DKO cells completely eliminate the surrounding cells.

In the laser ablation experiments, kymographs were generated using Fiji/ImageJ. The length of the junction-of-interest was measured before and after laser ablation, and divided by the mean junction length before laser ablation to yield the relative junction length. The recoil velocity was determined as the immediate length change of the junction-of-interest in response to laser ablation.

For quantification of the cell shape, cell outlines were segmented from the microscope images of the co-culture that were stained for cell–cell junction markers. The surrounding cells and *ZO-1*/*ZO-2* DKO cells were determined on the basis of the ZO-1 staining. The apical cell area of the cells was measured using Fiji/ImageJ. The aspect ratios of the cells were measured by fitting ellipsoids and taking the ratio of the major axis and the minor axis. The boundary cell angle was determined by measuring the relative angle between the clonal boundary and the major axis of the surrounding cell located at the boundary.

TAZ/YAP localization was quantified by counting the number of cells that exhibited distinct nuclear localization of TAZ/YAP and dividing them by the total number of cells. For co-culture experiments, the *ZO-1*/*ZO-2* DKO cells were distinguished from the surrounding cells based on ZO-1 staining.

Statistical tests (unpaired two-tailed *t*-tests or one-way ANOVA with Tukey or Bonferroni post-hoc testing) were performed and plots were generated using Excel (Microsoft) or R (R foundation).

## References

1. Varadarajan, S., Stephenson, R.E., and Miller, A.L. (2019). Multiscale dynamics of tight junction remodeling. J. Cell Sci. 132, jcs229286. 10.1242/jcs.229286.

2. Farquhar, M.G., and Palade, G.E. (1963). Junctional complexes in various epithelia. J. Cell Biol. 17, 375–412. 10.1083/jcb.17.2.375

3. Takeichi, M. (2014). Dynamic contacts: Rearranging adherens junctions to drive epithelial remodeling. Nat. Rev. Mol. Cell Biol. 15, 397–410. 10.1038/nrm3802

4. Otani, T., and Furuse, M. (2020). Tight junction structure and function revisited. Trends Cell Biol. 30, 805–817. 10.1016/j.tcb.2020.08.004.

5. Zihni, C., Mills, C., Matter, K., and Balda, M.S. (2016). Tight junctions: From simple barriers to multifunctional molecular gates. Nat. Rev. Mol. Cell Biol. 17, 564–580. 10.1038/nrm.2016.80

6. Umeda, K., Ikenouchi, J., Katahira-Tayama, S., Furuse, K., Sasaki, H., Nakayama, M., Matsui, T., Tsukita, S., Furuse, M., and Tsukita, S. (2006). ZO-1 and ZO-2 independently determine where claudins are polymerized in tight-junction strand formation. Cell 126, 741–754. 10.1016/j.cell.2006.06.043.

7. Ikenouchi, J., Umeda, K., Tsukita, S., Furuse, M., and Tsukita, S. (2007). Requirement of ZO-1 for the formation of belt-like adherens junctions during epithelial cell polarization. J. Cell Biol. 176, 779–786. 10.1083/jcb.200612080

8. Phua, D.C.Y., Xu, J., Ali, S.M., Boey, A., Gounko, N. V., and Hunziker, W. (2014). ZO-1 and ZO-2 are required for extra-embryonic endoderm integrity, primitive ectoderm survival and normal cavitation in embryoid bodies derived from mouse embryonic stem cells. PLoS One 9. e99532. 10.1371/journal.pone.0099532.

9. Otani, T., Nguyen, T.P., Tokuda, S., Sugihara, K., Sugawara, T., Furuse, K., Miura, T., Ebnet, K., and Furuse, M. (2019). Claudins and JAM-A coordinately regulate tight junction formation and epithelial polarity. J. Cell Biol. 218, 3372–3396. 10.1083/JCB.201812157.

10. Meng, W., and Takeichi, M. (2009). Adherens junction: molecular architecture and regulation. Cold Spring Harb. Perspect. Biol. 1, a002899. 10.1101/cshperspect.a002899.

11. Yonemura, S., Wada, Y., Watanabe, T., Nagafuchi, A., and Shibata, M. (2010). α-Catenin as a tension transducer that induces adherens junction development. Nat. Cell Biol. 12, 533–542. 10.1038/ncb2055.

12. Shen, L., Weber, C.R., Raleigh, D.R., Yu, D., and Turner, J.R. (2011). Tight junction pore and leak pathways: A dynamic duo. Annu. Rev. Physiol. 73, 283–309. 10.1146/annurev-physiol-012110-142150.

13. Nguyen, T.P., Otani, T., Tsutsumi, M., Kinoshita, N., Fujiwara, S., Nemoto, T., Fujimori, T., and Furuse, M. (2024). Tight junction membrane proteins regulate the mechanical resistance of the apical junctional complex. J. Cell Biol. 223, e202307104. 10.1083/jcb.202307104.

14. Madara, J.L. (1990). Maintenance of the macromolecular barrier at cell extrusion sites in intestinal epithelium: Physiological rearrangement of tight junctions. J. Membrane Biol. 116, 177–184. 10.1007/BF01868675.

15. Jinguji, Y., and Ishikawa, H. (1992). Electron microscopic observations on the maintenance of the tight junction during cell division in the epithelium of the mouse small intestine. Cell Struct. Function 17, 27–37. 10.1247/csf.17.27.

16. Baker, J., and Garrod, D. (1993). Epithelial cells retain junctions during mitosis. J. Cell Sci. 104, 415–425. 10.1242/jcs.104.2.415.

17. Guan, Y., Watson, A.J.M., Marchiando, A.M., Bradford, E., Shen, L., Turner, J.R., and Montrose, M.H. (2011). Redistribution of the tight junction protein ZO-1 during physiological shedding of mouse intestinal epithelial cells. Am. J. Physiol. -Cell Physiol. 300. C1404–1414. 10.1152/ajpcell.00270.2010.

18. Higashi, T., Arnold, T.R., Stephenson, R.E., Dinshaw, K.M., and Miller, A.L. (2016). Maintenance of the epithelial barrier and remodeling of cell-cell junctions during cytokinesis. Curr. Biol. 26, 1829–1842. 10.1016/j.cub.2016.05.036.

19. Stephenson, R.E., Higashi, T., Erofeev, I.S., Arnold, T.R., Leda, M., Goryachev, A.B., and Miller, A.L. (2019). Rho flares repair local tight junction leaks. Dev. Cell 48, 445–459.e5. 10.1016/j.devcel.2019.01.016.

20. Higashi, T., Saito, A.C., Fukazawa, Y., Furuse, M., Higashi, A.Y., Ono, M., and Chiba, H. (2023). EpCAM proteolysis and release of complexed claudin-7 repair and maintain the tight junction barrier. J. Cell Biol. 222. e202204079. 10.1083/jcb.202204079.

21. Cho, Y., Taniguchi, A., Kubo, A., and Ikenouchi, J. (2024). Rho-ROCK liberates sequestered claudin for rapid de novo tight junction formation. bioRxiv. 10.1101/2024.09.09.612007.

22. Oda, Y., Takahashi, C., Harada, S., Nakamura, S., Sun, D., Kiso, K., Urata, Y., Miyachi, H., Fujiyoshi, Y., Honigmann, A., et al. (2021). Discovery of anti-inflammatory physiological peptides that promote tissue repair by reinforcing epithelial barrier formation. Science Advances 7, eabj6895. 10.1126/sciadv.abj6895.

23. Clavería, C., and Torres, M. (2016). Cell competition: Mechanisms and physiological roles. Annu. Rev. Cell Dev. Biol. 32, 411–439. 10.1146/annurev-cellbio-111315-125142.

24. van Neerven, S.M., and Vermeulen, L. (2023). Cell competition in development, homeostasis and cancer. Nat. Rev. Mol. Cell Biol. 24, 221–236. 10.1038/s41580-022-00538-y.

25. Morata, G., and Ripoll, P. (1975). Minutes: Mutants of Drosophila autonomously affecting cell division rate. Dev. Biol. 42, 211–221. 10.1016/0012-1606(75)90330-9.

26. Hogan, C., Dupré-Crochet, S., Norman, M., Kajita, M., Zimmermann, C., Pelling, A.E., Piddini, E., Baena-López, L.A., Vincent, J.P., Itoh, Y., et al. (2009). Characterization of the interface between normal and transformed epithelial cells. Nat. Cell Biol. 11, 460–467. 10.1038/ncb1853.

27. Igaki, T., Pastor-Pareja, J.C., Aonuma, H., Miura, M., and Xu, T. (2009). Intrinsic tumor suppression and epithelial maintenance by endocytic activation of Eiger/TNF signaling in Drosophila. Dev. Cell 16, 458–465. 10.1016/j.devcel.2009.01.002.

28. Clavería, C., Giovinazzo, G., Sierra, R., and Torres, M. (2013). Myc-driven endogenous cell competition in the early mammalian embryo. Nature 500, 39–44. 10.1038/nature12389.

29. Sancho, M., Di-Gregorio, A., George, N., Pozzi, S., Sánchez, J.M., Pernaute, B., and Rodríguez, T.A. (2013). Competitive interactions eliminate unfit embryonic stem cells at the onset of differentiation. Dev. Cell 26, 19–30. 10.1016/j.devcel.2013.06.012.

30. Akieda, Y., Ogamino, S., Furuie, H., Ishitani, S., Akiyoshi, R., Nogami, J., Masuda, T., Shimizu, N., Ohkawa, Y., and Ishitani, T. (2019). Cell competition corrects noisy Wnt morphogen gradients to achieve robust patterning in the zebrafish embryo. Nat. Commun. 10, 4710. 10.1038/s41467-019-12609-4.

31. Ellis, S.J., Gomez, N.C., Levorse, J., Mertz, A.F., Ge, Y., and Fuchs, E. (2019). Distinct modes of cell competition shape mammalian tissue morphogenesis. Nature 569, 497–502. 10.1038/s41586-019-1199-y.

32. Hashimoto, M., and Sasaki, H. (2019). Epiblast formation by TEAD-YAP-dependent expression of pluripotency factors and competitive elimination of unspecified cells. Dev. Cell 50, 139–154.e5. 10.1016/j.devcel.2019.05.024.

33. Liu, N., Matsumura, H., Kato, T., Ichinose, S., Takada, A., Namiki, T., Asakawa, K., Morinaga, H., Mohri, Y., De Arcangelis, A., et al. (2019). Stem cell competition orchestrates skin homeostasis and ageing. Nature 568, 344–350. 10.1038/s41586-019-1085-7.

34. Moya, I.M., Castaldo, S.A., Van Den Mooter, L., Soheily, S., Sansores-Garcia, L., Jacobs, J., Mannaerts, I., Xie, J., Verboven, E., Hillen, H., et al. (2019). Peritumoral activation of the Hippo pathway effectors YAP and TAZ suppresses liver cancer in mice. Science 366, 1029–1034. 10.1126/science.aaw9886.

35. Czabotar, P., Lessene, G., Strasser, A., and Adams, J.M. (2013) Control of apoptosis by the BCL-2 protein family: implications for physiology and therapy. Nat. Rev. Mol. Cell Biol. 15, 49–63. 10.1038/nrm3722.

36. Amano, M., Nakayama, M., and Kaibuchi, K. (2010). Rho-kinase/ROCK: A key regulator of the cytoskeleton and cell polarity. Cytoskeleton 67, 545–554. 10.1002/cm.20472.

37. Dupont, S., Morsut, L., Aragona, M., Enzo, E., Giulitti, S., Cordenonsi, M., Zanconato, F., Le Digabel, J., Forcato, M., Bicciato, S., et al. (2011). Role of YAP/TAZ in mechanotransduction. Nature 474, 179–184. 10.1038/nature10137.

38. Domínguez-Calderón, A., Ávila-Flores, A., Ponce, A., López-Bayghen, E., Calderón-Salinas, J.V., Reyes, J.L., Chávez-Munguía, B., Segovia, J., Angulo, C., Ramírez, L., et al. (2016). ZO-2 silencing induces renal hypertrophy through a cell cycle mechanism and the activation of YAP and the mTOR pathway. Mol. Biol. Cell 27, 1581–1595. 10.1091/mbc.E15-08-0598.

39. Haas, A.J., Karakus, M., Zihni, C., Balda, M.S., and Matter, K. (2024). ZO-1 regulates Hippo-independent YAP activity and cell proliferation via a GEF-H1- and TBK1-regulated signalling network. Cells 13. 10.3390/cells13070640.

40. Kwon, H., Kim, J., and Jho, E. (2022). Role of the Hippo pathway and mechanisms for controlling cellular localization of YAP/TAZ. FEBS Journal 289, 5798–5818. 10.1111/febs.16091.

41. Chumki, S.A., van den Goor, L.M., Hall, B.N., and Miller, A.L. (2022). p115RhoGEF activates RhoA to support tight junction maintenance and remodeling. Mol. Biol. Cell 33. 10.1091/mbc.E22-06-0205.

42. Valon, L., Matamoro-Vidal, A., Villars, A., and Levayer, R. (2024). Interfacial tension and growth both contribute to mechanical cell competition. Submitted.

43. Eisenhoffer, G.T., Loftus, P.D., Yoshigi, M., Otsuna, H., Chien, C. Bin, Morcos, P.A., and Rosenblatt, J. (2012). Crowding induces live cell extrusion to maintain homeostatic cell numbers in epithelia. Nature 484, 546–549. 10.1038/nature10999.

44. Wagstaff, L., Goschorska, M., Kozyrska, K., Duclos, G., Kucinski, I., Chessel, A., Hampton-O’Neil, L., Bradshaw, C.R., Allen, G.E., Rawlins, E.L., et al. (2016). Mechanical cell competition kills cells via induction of lethal p53 levels. Nat. Commun. 7. 10.1038/ncomms11373.

45. Otani, T., Ichii, T., Aono, S., and Takeichi, M. (2006). Cdc42 GEF Tuba regulates the junctional configuration of simple epithelial cells. J. Cell Biol. 175, 135–146. 10.1083/jcb.200605012.

46. Van Itallie, C.M., Fanning, A.S., Bridges, A., and Anderson, J.M. (2009). ZO-1 Stabilizes the Tight Junction Solute Barrier through Coupling to the Perijunctional Cytoskeleton. Mol. Biol. Cell 20, 3930–3940. 10.1091/mbc.E09.

47. Fanning, A.S., Van Itallie, C.M., and Anderson, J.M. (2012). Zonula occludens-1 and -2 regulate apical cell structure and the zonula adherens cytoskeleton in polarized epithelia. Mol. Biol. Cell 23, 577–590. 10.1091/mbc.E11-09-0791.

48. Choi, W., Acharya, B.R., Peyret, G., Fardin, M.A., Mège, R.M., Ladoux, B., Yap, A.S., Fanning, A.S., and Peifer, M. (2016). Remodeling the zonula adherens in response to tension and the role of afadin in this response. J. Cell Biol. 213, 243–260. 10.1083/jcb.201506115.

49. Bement, W.M., Forscher, P., and Mooseker, M.S. (1993). A novel cytoskeletal structure involved in purse string wound closure and cell polarity maintenance. J. Cell Biol. 121, 565–578. 10.1038/jcb.121.3.565.

50. Brock, J., Midwinter, K., Lewis, J., and Martin, P. (1996). Healing of incisional wounds in the embryonic chick wing bud: Characterization of the actin purse-string and demonstration of a requirement for Rho activation. J. Cell Biol. 135, 1097–1107. 10.1083/jcb.135.4.1097.

51. Florian, P., Schöneberg, T., Schulzke, J.D., Fromm, M., and Gitter, A.H. (2002). Single-cell epithelial defects close rapidly by an actinomyosin purse string mechanism with functional tight junctions. J. Physiol. 545, 485–499. 10.1113/jphysiol.2002.031161.

52. Rosenblatt, J., Raff, M.C., and Cramer, L.P. (2001). An epithelial cell destined for apoptosis signals its neighbors to extrude it by an actin- and myosin-dependent mechanism. Curr. Biol. 11, 1847–1857. 10.1016/S0960-9822(01)00587-5.

53. Young, P.E., Richman, A.M., Ketchum, A.S., and Kiehart, D.P. (1993). Morphogenesis in Drosophila requires nonmuscle myosin heavy chain function. Genes Dev. 7, 29–41. 10.1101/gad.7.1.29.

54. Kiehart, D.P., Galbraith, C.G., Edwards, K.A., Rickoll, W.L., and Montague, R.A. (2000). Multiple Forces Contribute to Cell Sheet Morphogenesis for Dorsal Closure in Drosophila. J. Cell Biol. 149, 471–490. 10.1083/jcb.149.2.471.

55. Hutson, M.S., Tokutake, Y., Chang, M.-S., Bloor, J.W., Venakides, S., Kiehart, D.P., and Edwards, G.S. (2003). Forces for Morphogenesis Investigated with Laser Microsurgery and Quantitative Modeling. Science 300, 145–149. 10.1126/science.1079552.

56. Duszyc, K., Gomez, G.A., Lagendijk, A.K., Yau, M.K., Nanavati, B.N., Gliddon, B.L., Hall, T.E., Verma, S., Hogan, B.M., Pitson, S.M., et al. (2021). Mechanotransduction activates RhoA in the neighbors of apoptotic epithelial cells to engage apical extrusion. Curr. Biol. 31, 1326–1336.e5. 10.1016/j.cub.2021.01.003.

57. Yu, H.H., and Zallen, J.A. (2020). Abl and Canoe/Afadin mediate mechanotransduction at tricellular junctions. Science 370, eaba5528. 10.1126/science.aba5528.

58. Vachharajani, V.T., DeJong, M.P., and Dunn, A.R. (2023). PDZ Domains from the Junctional Proteins Afadin and ZO-1 Act as Mechanosensors. bioRxiv. 10.1101/2023.09.24.559210.

59. Bonello, T.T., Perez-Vale, K.Z., Sumigray, K.D., and Peifer, M. (2018). Rap1 acts via multiple mechanisms to position canoe and adherens junctions and mediate apical-basal polarity establishment. Development 145, dev157941. 10.1242/dev.157941. 58

60. Schmidt, A., Lv, Z., and Großhans, J. (2018). Elmo and sponge specify subapical restriction of canoe and formation of the subapical domain in early drosophila embryos. Development 145, dev157909. 10.1242/dev.157909.

61. Perez-Vale, K.Z., Yow, K.D., Johnson, R.I., Byrnes, A.E., Finegan, T.M., Slep, K.C., and Peifer, M. (2021). Multivalent interactions make adherens junction– cytoskeletal linkage robust during morphogenesis. J. Cell Biol. 220, e202104087. 10.1083/jcb.202104087.

62. Gurley, N.J., Szymanski, R.A., Dowen, R.H., Butcher, T.A., Ishiyama, N., and Peifer, M. (2023). Exploring the evolution and function of Canoe’s intrinsically disordered region in linking cell-cell junctions to the cytoskeleton during embryonic morphogenesis. PLoS One 18, e0289224. 10.1371/journal.pone.0289224.

63. McParland, E.D., Gurley, N.J., Wolfsberg, L.R., Butcher, T.A., Bhattarai, A., Jensen, C.C., Johnson, R.I., Slep, K.C., and Peifer, M. (2024). The dual Ras Association (RA) domains of Drosophila Canoe have differential roles in linking cell junctions to the cytoskeleton during morphogenesis. J. Cell Sci. jcs.263546. 10.1242/jcs.263546.

64. McParland, E.D., Butcher, T.A., Gurley, N.J., Johnson, R.I., Slep, K.C., and Peifer, M. (2024). The Dilute domain in Canoe is not essential for linking cell junctions to the cytoskeleton but supports morphogenesis robustness. J. Cell Sci. 137. 10.1242/jcs.261734.

65. Kuno, S., Nakamura, R., Otani, T., and Togashi, H. (2024). Multivalent Afadin Interaction Promotes IDR-Mediated Condensate Formation and Junctional Separation of Epithelial Cells. bioRxiv. 10.1101/2024.04.26.591237.

66. Sawyer, J.K., Harris, N.J., Slep, K.C., Gaul, U., and Peifer, M. (2009). The Drosophila afadin homologue Canoe regulates linkage of the actin cytoskeleton to adherens junctions during apical constriction. J. Cell Biol. 186, 57–73. 10.1083/jcb.200904001.

67. Sawyer, J.K., Choi, W., Jung, K.-C., He, L., Harris, N.J., and Peifer, M. (2011). A contractile actomyosin network linked to adherens junctions by Canoe/afadin helps drive convergent extension. Mol. Biol. Cell 22, 2491–2508. 10.1091/E11-05-0411.

68. Mamada, H., Sato, T., Ota, M., and Sasaki, H. (2015). Cell competition in mouse NIH3T3 embryonic fibroblasts is controlled by the activity of Tead family proteins and Myc. J. Cell Sci. 128, 790–803. 10.1242/jcs.163675.

69. Elbediwy, A., Vincent-Mistiaen, Z.I., and Thompson, B.J. (2016). YAP and TAZ in epithelial stem cells: A sensor for cell polarity, mechanical forces and tissue damage. BioEssays 38, 644–653. 10.1002/bies.201600037.

70. Lee, M.J., Byun, M.R., Furutani-Seiki, M., Hong, J.H., and Jung, H.S. (2014). YAP and TAZ regulate skin wound healing. J. Invest. Dermatol. 134, 518–525. 10.1038/jid.2013.339.

71. Cartagena-Rivera, A.X., Van Itallie, C.M., Anderson, J.M., and Chadwick, R.S. (2017). Apical surface supracellular mechanical properties in polarized epithelium using noninvasive acoustic force spectroscopy. Nat. Commun. 8, 1030. 10.1038/s41467-017-01145-8.

72. Martin, T.A., and Jiang, W.G. (2009). Loss of tight junction barrier function and its role in cancer metastasis. Biochim. Biophys. Acta. 1788, 872–891. 10.1016/j.bbamem.2008.11.005.

73. Bastounis, E.E., Serrano-Alcalde, F., Radhakrishnan, P., Engström, P., Gómez-Benito, M.J., Oswald, M.S., Yeh, Y.T., Smith, J.G., Welch, M.D., García-Aznar, J.M., et al. (2021). Mechanical competition triggered by innate immune signaling drives the collective extrusion of bacterially infected epithelial cells. Dev. Cell 56, 443–460.e11. 10.1016/j.devcel.2021.01.012.

74. Schoenenberger, C.A., Zuk, A., Kendall, D., and Matlin, K.S. (1991). Multilayering and Loss of Apical Polarity in MDCK Cells Transformed with Viral K-ras. J. Cell Biol. 112, 873–889. 10.1083/jcb.112.5.873.

75. Matsuyoshi, N., Hamaguchi, M., Taniguchi, S., Nagafuchi, A., Tsukita, S., and Takeichi, M. (1992). Cadherin-Mediated Cell-Cell Adhesion Is Perturbed by v-src Tyrosine Phosphorylation m Metastatic Fibroblasts. J. Cell Biol. 118, 703–714. 10.1083/jcb.118.3.703.

76. Behrens, J., Vakaet, L., Friis, R., Winterhager, E., Van Roy, F., Mareel, M.M., and Birchmeier, W. (1993). Loss of Epithelial Differentiation and Gain of Invasiveness Correlates with Tyrosine Phosphorylation of the E-Cadherin/beta-Catenin Complex in Cells Transformed with a Temperature-sensitive v-SRC Gene. J. Cell Biol. 120, 757–766. 10.1083/jcb.120.3.757.

77. Kinch, M.S., Clark, G.J., Der, C.J., and Burridge, K. (1995). Tyrosine Phosphorylation Regulates the Adhesions of Ras-transformed Breast Epithelia. J. Cell Biol. 130, 461–471. 10.1083/jcb.130.2.461.

78. Kajita, M., Hogan, C., Harris, A.R., Dupre-Crochet, S., Itasaki, N., Kawakami, K., Charras, G., Tada, M., and Fujita, Y. (2010). Interaction with surrounding normal epithelial cells influences signalling pathways and behaviour of Src-transformed cells. J. Cell Sci. 123, 171–180. 10.1242/jcs.057976.

79. Wang, F., Graham, W., Vallen Wang, Y., Witkowski, E.D., Schwarz, B.T., and Turner, J.R. (2005). Interferon-and Tumor Necrosis Factor-Synergize to Induce Intestinal Epithelial Barrier Dysfunction by Up-Regulating Myosin Light Chain Kinase Expression. Am. J. Pathol. 166, 409–419. 10.1016/s0002-9440(10)62264-x.

80. Marchiando, A.M., Shen, L., Graham, W.V., Edelblum, K.L., Duckworth, C.A., Guan, Y., Montrose, M.H., Turner, J.R., and Watson, A.J.M. (2011). The Epithelial Barrier Is Maintained by In Vivo Tight Junction Expansion During Pathologic Intestinal Epithelial Shedding. Gastroenterology 140, 1208–1218.e2. 10.1053/j.gastro.2011.01.004.

81. Bagley, D.C., Russell, T., Ortiz-Zapater, E., Stinson, S., Fox, K., Redd, P.F., Joseph, M., Deering-Rice, C., Reilly, C., Parsons, M., et al. (2024). Bronchoconstriction damages airway epithelia by crowding-induced excess cell extrusion. Science 384, 66–73. 10.1126/science.adk2758.

82. Richardson, J.C.W., Scalera, V., and Simmons, N.L. (1981). Identification of two strains of MDCK cells which resemble separate nephron tubule segments. Biochim. Biophys. Acta - Gen. Subj. 673, 26–36. 10.1016/0304-4165(81)90307-X.

83. Reichmann, E., Schwarz, H., Deiner, E.M., Leitner, I., Eilers, M., Berger, J., Busslinger, M., and Beug, H. (1992). Activation of an inducible c-FosER fusion protein causes loss of epithelial polarity and triggers epithelial-fibroblastoid cell conversion. Cell 71, 1103–1116. 10.1016/S0092-8674(05)80060-1.

84. Ichii, T., and Takeichi, M. (2007). p120-catenin regulates microtubule dynamics and cell migration in a cadherin-independent manner. Genes Cells 12, 827– 839. 10.1111/j.1365-2443.2007.01095.x.

85. Wang, N.S., Unkila, M.T., Reineks, E.Z., and Distelhorst, C.W. (2001). Transient Expression of Wild-type or Mitochondrially Targeted Bcl-2 Induces Apoptosis, whereas Transient Expression of Endoplasmic Reticulum-targeted Bcl-2 Is Protective against Bax-induced Cell Death. J. Biol. Chem. 276, 44117–44128. 10.1074/jbc.M101958200.

86. Riedl, J., Crevenna, A.H., Kessenbrock, K., Yu, J.H., Neukirchen, D., Bista, M., Bradke, F., Jenne, D., Holak, T.A., Werb, Z., et al. (2008). Lifeact: a versatile marker to visualize F-actin. Nat. Methods 5, 605–607. 10.1038/nmeth.1220.

87. Asano, S., Hamao, K., and Hosoya, H. (2009). Direct evidence for roles of phosphorylated regulatory light chain of myosin II in furrow ingression during cytokinesis in HeLa cells. Genes to Cells 14, 555–568. 10.1111/j.1365-2443.2009.01288.x.

88. Hirate, Y., Hirahara, S., Inoue, K.I., Suzuki, A., Alarcon, V.B., Akimoto, K., Hirai, T., Hara, T., Adachi, M., Chida, K., et al. (2013). Polarity-dependent distribution of angiomotin localizes hippo signaling in preimplantation embryos. Curr. Biol. 23, 1181–1194. 10.1016/j.cub.2013.05.014.

89. Uehata, M., Ishizaki, T., Satoh, H., Ono, T., Kawahara, T., Morishita, T., Tamakawa, H., Yamagami, K., Inui, J., Maekawa, M., et al. (1997). Calcium sensitization of smooth muscle mediated by a Rho-associated protein kinase in hypertension. Nature 389, 990–994. 10.1038/40187.

90. Straight, A.F., Cheung, A., Limouze, J., Chen, I., Westwood, N.J., Sellers, J.R., and Mitchison, T.J. (2003). Dissecting Temporal and Spatial Control of Cytokinesis with a Myosin II Inhibitor. Science. 299, 1743–1747. 10.1126/science.1081412.

91. Itoh, M., Yonemura, S., Nagafuchi, A., Tsukita, S., and Tsukita, S. (1991). A 220-kD undercoat-constitutive protein: its specific localization at cadherin-based cell-cell adhesion sites. J. Cell Biol. 115, 1449–1462. 10.1083/jcb.115.5.1449.

92. Stevenson, B.R., Siliciano, J.D., Mooseker, M.S., and Goodenough, D.A. (1986). Identification of ZO-1: a high molecular weight polypeptide associated with the tight junction (zonula occludens) in a variety of epithelia. J. Cell Biol. 103, 755–766. 10.1083/jcb.103.3.755.

93. Nagafuchi, A., and Tsukita, S. (1994). The Loss of the Expression of α Catenin, the 102 kD Cadherin Associated Protein, in Central Nervous Tissues during Development. Dev. Growth Differ. 36, 59–71. 10.1111/j.1440-169X.1994.00059.x.

